# Methionine modulates the metabolic-epigenetic axis as immunotherapeutic in tuberculosis

**DOI:** 10.1101/2025.10.07.680834

**Authors:** Nidhi Yadav, Ashish Gupta, Satya Ranjan Sahoo, Suchitra Jena, Nainy Goel, Mothe Sravya, Tundup Namgail, Nupur Sharma, Someshwar Nath Jha, Bhishma Narayan Panda, Nimesh Gupta, Jaswinder Singh Maras, Amit Kumar Pandey, Shyam Kumar Masakapalli, Debasis Dash, Ranjan Kumar Nanda

**Affiliations:** Translational Health Group, International Centre for Genetic Engineering and Biotechnology, New Delhi- 110067, India; School of Life Sciences, Sambalpur University, Jyoti Vihar, Burla, Odisha-768019, India; Institute of Life Sciences, Nalco Nagar Rd, NALCO Square, Bhubaneswar, Odisha-751023, India; Department of Molecular and Cellular Medicine, Institute of Liver and Biliary Sciences, New Delhi-110070, India; Vaccine Immunology Laboratory, National Institute of Immunology, New Delhi-110070, India; Translational Health Science and Technology Institute, NCR Biotech Science Cluster, 3rd Milestone, Faridabad, Haryana-121001, India; School of Biosciences and Bioengineering, Indian Institute of Technology Mandi, Kamand-175075, India

**Keywords:** Tuberculosis, Bone marrow-derived macrophages, Metabolomics, Methionine metabolism, Polyamine synthesis, Nucleotide salvage

## Abstract

The methionine metabolism is central to epigenetic reprogramming to produce pro-inflammatory cytokines. Disruptions in methionine metabolism contribute to complex disorders, providing an important target for nutrient interventions. Here, *Mycobacterium tuberculosis* (Mtb) H37Rv-infected C57BL/6 mice showed functional heterogeneity between alveolar and non-alveolar macrophage (AMs/Non-AMs) with major metabolic reprogramming in Non-AMs. Global metabolite and proteome analysis of Mtb-infected bone marrow-derived macrophages (BMDMs) showed a diversion of flux from methionine metabolism toward increased nucleotide salvage and glutathione (GSH) production. Carbon units of ^13^C_5_-methionine via isotopomer analysis contributed to polyamine synthesis, fuelling the nucleotide salvage node in Mtb-infected BMDMs. Methionine supplementation increased mycobacterial clearance in C57BL/6 mice and in BMDMs by promoting a pro-inflammatory response, as evident by increased IL-1β and IFN-γ. Increased IL-1β production is mainly contributed to by increased H3K4 trimethylation in macrophages. These findings reveal an important strategy by which methionine supplementation in macrophages and mice remodels host metabolism to enhance acute pro-inflammatory responses and exploit dietary supplementation to improve nutritional immunity in TB.

**Highlights:**

- *Mycobacterium tuberculosis* (Mtb) infection decreases intracellular methionine levels and increases purine levels in primary macrophages.
- Methionine carbon units contribute to nucleotide salvage, and their restriction lowers IL-1 response and mycobacterial clearance.
- *Ex vivo* methionine supplementation in Mtb-infected macrophages increases H3K4me3 at the *Il-1b* gene promoter and gene body, leading to an increased pro-inflammatory response and improved mycobacterial clearance.
- *In vivo* methionine supplementation improved pro-inflammatory response in lungs, spleen and bone marrow, leading to accelerated mycobacterial clearance upon Mtb-H37Rv infection.

**Graphical abstract:** 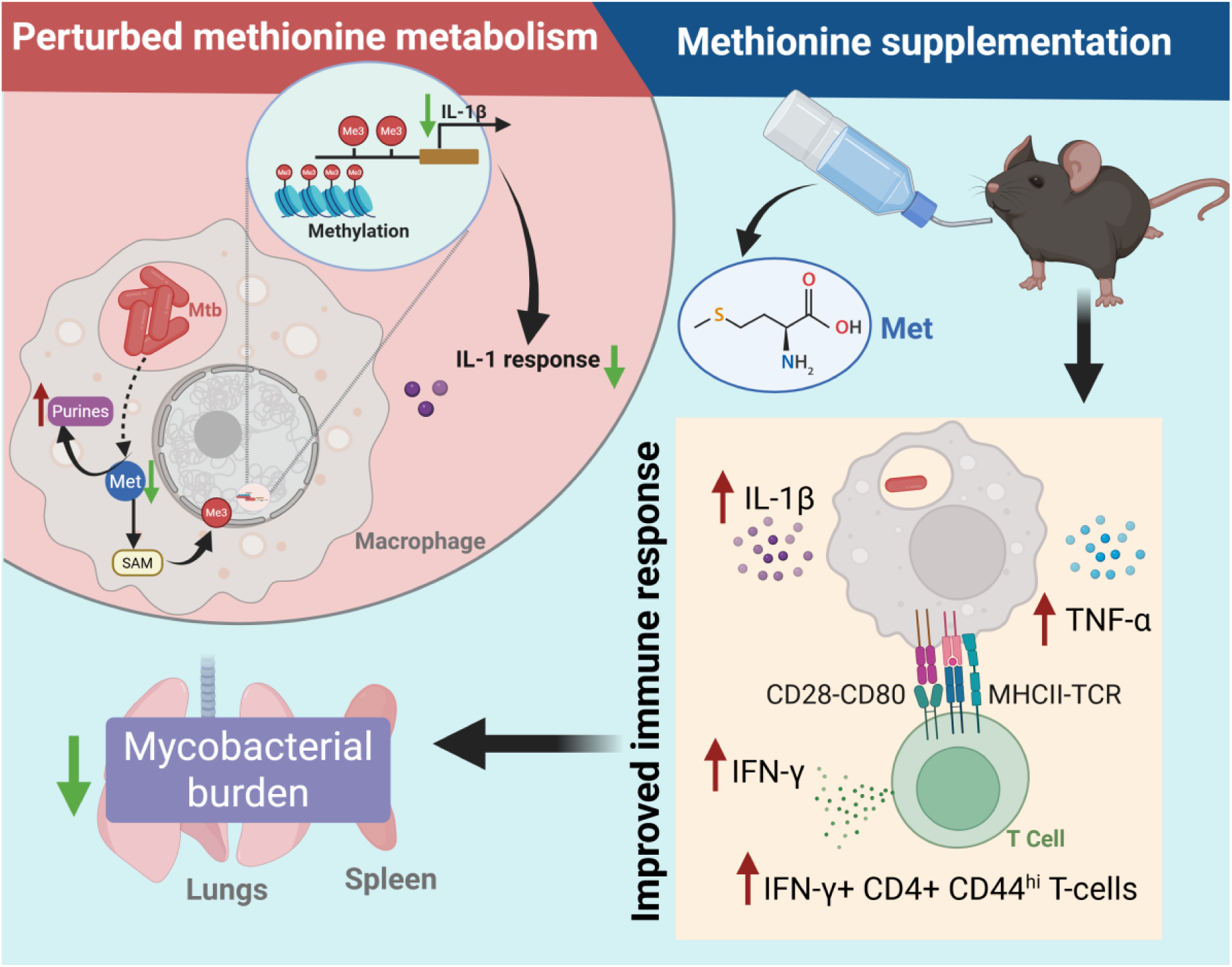

## Introduction

*Mycobacterium tuberculosis* (Mtb), the causative organism of tuberculosis (TB), transmits from the aerosol droplets generated during the cough of a patient to the lungs of a new host. The lung alveolar macrophages are among the first responders to incoming Mtb, and upon infection, lead to metabolic reprogramming (Borah Slater et al., 2023), and the release of chemokines and cytokines, which increase neutrophil recruitment. (Madden et al., 2022) From the airway epithelium, Mtb-infected alveolar macrophages migrate into the lung interstitium. (Cohen et al., 2018) Energy generation in the alveolar and interstitial macrophages primarily depends on oxidative phosphorylation and glycolysis, respectively. Mtb-infected alveolar macrophages show M2 polarisation, with increased glutamine and lipid uptake, and are associated with an anti-inflammatory response, as evident by higher IL-10 production, which provides a favourable niche for Mtb replication, leading to a higher mycobacterial burden.

However, the interstitial macrophages show an M1 phenotype, associated with increased glycolysis, glutaminolysis and pro-inflammatory cytokine production, leading to increased bacterial clearance. (Wculek et al., 2022) Central carbon and arginine metabolism contribute to these differences in the polarisation-associated phenotypes in the macrophages. (Rath et al., 2014)(Rath et al., 2014)(Tiwari et al., 2018)(Wculek et al., 2022) Pro-inflammatory macrophages have higher iNOS expression with more nitric oxide formation for anti- microbial activity, whereas the anti-inflammatory phenotype has higher arginase expression, leading to higher ornithine and polyamine synthesis. One-carbon metabolism has emerged as a key metabolic node in proliferating immune cells, with a network of interconnected pathways for the transfer of one-carbon units. (Locasale, 2013) Methionine metabolism and the Folate cycle are the active carrier nodes for one-carbon units needed for nucleotide salvage and de novo nucleotide synthesis, respectively. The methionine cycle contributes to S-adenosyl methionine (SAM) production, which donates a methyl group during methylation and is also linked to glutathione production, a major cellular antioxidant. One-carbon units from the methionine cycle are reported to have immunomodulatory effects in LPS-induced inflammation. LPS-stimulated macrophages showed Histone H3 lysine 36 trimethylation by SAM, resulting in increased IL-1β production. (Yu et al., 2019) IL-1 responses are the major contributors to early-phase protective immunity against Mtb. (Juffermans et al., 2000)(Jayaraman et al., 2013) Macrophages, upon encountering Mtb, produce IL-1β and control the infection. (Yamada et al., 2000) However, limited studies elucidated the role of one-carbon metabolism and its impact on IL-1β production in Mtb-infected macrophages.

In this study, we investigated the dynamics of various interconnected nodes of methionine metabolism in Mtb-H37Rv-infected macrophages and in Mtb-H37Rv-infected mice. Our earlier report on single-cell RNA sequencing (scRNAseq) data analysis showed that alveolar macrophages (AMs) from Mtb-infected mice exhibited an impaired glycolytic switch and dysregulation of one-carbon metabolism compared with non-AMs. (Chaudhary et al., 2025) Primary bone marrow-derived macrophages (BMDMs) at 24 hours post-Mtb infection showed a significantly higher mycobacterial burden, lower levels of methionine, serine, and polyamines, and an upregulated nucleotide salvage and glutathione pathway. Interestingly, methionine supplementation (1 mM) in Mtb H37Rv-infected macrophages improved cell viability and significantly increased IL-1β production by increasing chromatin occupancy of the *Il-1b* gene via H3K4me3, resulting in a decreased mycobacterial burden. Supplementation of methionine in the drinking water of C57BL/6 mice improved lung mycobacterial clearance. Continuous methionine supplementation (∼2.5%), 2 weeks prior to infection and continued for 3 weeks post-infection, enhanced lung Mtb clearance with higher IL-1β and IFN-γ production. Methionine supplementation for 2 weeks, followed by discontinuation a day prior to infection, also showed an immunoprophylactic effect by priming innate immune cells for a more robust pro-inflammatory response and activating T cells, leading to a Mtb-specific IFN-γ response in the spleen and improving pathogen clearance. The findings of this study highlight the importance of methionine metabolism in TB and its importance as a target for host-directed therapeutics to ameliorate its pathology.

## Methodology

### Ethical statement

The experimental procedures adopted in this study were approved by the Institute Biosafety Committee and the Institute Animal Ethics Committee of the International Centre for Genetic Engineering and Biotechnology, New Delhi (ICGEB/IAEC/25092023/39.6, extension dated 17/06/2025).

### scRNAseq experiment

Raw data of the scRNAseq experiment, performed earlier in the laboratory, were analysed to compare the transcriptomic differences in the alveolar macrophages between Mtb-H37Rv aerosol-infected C57BL/6 mice (TB) and healthy controls (HC). The detailed experimental strategy adopted in this scRNAseq study was explained in the earlier report. (Chaudhary et al., 2025) Briefly, male C57BL/6 mice (16-18 weeks) were infected with a low aerosol dose (100-120 CFU) of animal-passaged Mtb H37Rv using a Madison chamber in the tuberculosis aerosol challenge BSL-III facility (TACF) at the host institution. At 21 days post-infection (dpi), lungs were harvested and, single-cell suspensions were prepared by enzymatic digestion. After enumerating the cells using Countess 3 (Thermo Fisher Scientific, USA), viability was determined by trypan blue staining, and the cells were centrifuged. To the cell pellet, 94 μl of chilled PBS supplemented with FBS (10%) was added, and after adding TruStain FcX (0.5 μl) and True-Stain Monocyte Blocker (5 μl), the cells were incubated for 10 minutes at 4°C. To respective samples, BioLegend TotalSeq^TM^-B hashtags (0.5 μl) were added and incubated for 30 minutes at 4°C. Samples were diluted, and after centrifugation, the cells were incubated with diluted live/dead stain and FACS antibody mix (BioLegend). After dilution and centrifugation, the cells were resuspended in PBS supplemented with FBS for flow sorting using the BD FACS ARIA flow sorter present in the TACF.

Briefly, flow-sorted lung CD3+ and CD11c+ cells were mixed in a ratio of 2:1 and fixed using the supplied buffer (1 ml, 10x Genomics PN-2000517) for 24 hours at 4°C. After centrifuging, the cells were resuspended in quenching solution (1 ml, 10x Genomics, USA PN-2000516) and to it, Glycerol (50%, 275 µl, G5516, Millipore Sigma) and enhancer (10x Genomics, PN-2000482, 100 µl) were added and used for fixed single-cell RNA sequencing using 10x Genomics single-cell gene expression flex kit. Following the manufacturer’s instructions, probe hybridisation, GEM generation, barcoding and library preparation were performed. After checking the quality control of the prepared library using TapeStation (Agilent Technologies, USA), the gene expression (GEX) and cell surface protein (CSP) libraries were sequenced using PE150 chemistry on an SP100 flow cell at a commercial facility using the NovaSeq 6000 sequencing system (Illumina, USA). The sequencing depth was 10,000 read pairs/cell for GEX libraries and 5,000 read pairs/cell for CSP libraries. A 10% PhiX spike was used during library loading for the sequencing of these libraries. From the four study groups used in the original experiment, only the data sets from Mtb-infected and healthy controls were used for data analysis with standard pipelines.

### scRNAseq data analysis

Raw sequencing files from the NovaSeq6000 platform were processed using the Cell Ranger v7.2 multi-pipeline from 10x Genomics. Both raw and processed single-cell RNA-seq data are available in the Gene Expression Omnibus (GEO) under accession number GSE280089. Filtered gene expression and antibody capture matrices (filtered_feature_bc_matrix) were loaded using the Seurat R Package (v5.0.2). HTO assays were normalized using Centred Log-Ratio normalization, and demultiplexing of cells labelled with hashtag oligonucleotides (HTOs) was performed with the HTODemux function (positive quantile = 0.99). Singlet cells were retained and labelled as either “Tuberculosis; TB” or “Healthy control; HC”. Quality control was performed by measuring mitochondrial RNA percentage (percent.mt), number of genes detected per cell (nFeature_RNA), and total RNA counts per cell (nCount_RNA). For the TB sample, cells were retained if they had 200–5,000 detected genes, 600–15,000 total counts, and <5% mitochondrial content. For HC, the filtering criteria were 400–4200 detected genes, 700–10,000 total counts, and <5% mitochondrial content. Filtered datasets were log-normalised, and the top 2000 highly variable genes were identified using the Variance Stabilizing Transformation (VST) method. Datasets were integrated using Seurat’s anchor-based workflow to align shared cell populations across TB and HC samples. This enabled direct comparison of biological differences between conditions within a unified embedding space. First, integration anchors were identified via canonical correlation analysis (CCA) using the FindIntegrationAnchors function, based on the top 30 principal components. Genes common to both datasets were selected via gene name interactions and used as features for integration. The datasets were then integrated using IntegrateData with these common features. The integrated dataset was scaled, and dimensionality reduction was performed using principal component analysis (PCA), followed by Uniform Manifold Approximation and Projection (UMAP) for visualisation. Clusters were identified using the Louvain algorithm with a resolution of 0.7. PCA and UMAP plots were generated to assess integration quality and sample overlap. To annotate cell types, we employed the ScType pipeline using manually curated marker gene sets. (Ianevski et al., 2022) Marker enrichment scores were computed at the single-cell level using the sctype_score function on the scaled expression matrix. The top-scoring cell type per cluster was assigned as the annotation label. These labels were visualized on UMAP plots, both combined and split by condition. Marker genes were used for validation and visualization via DotPlot.

For pseudobulking and Differential gene expression analysis, specific cell populations were subsetted from the integrated Seurat object, based on cell type annotations obtained via Sctype. Pseudobulking is required for scRNA-seq differential expression analysis with DESeq2, as our scRNA-seq dataset included four biological replicates per group (TB and HC) and was multiplexed using cell-surface protein barcodes. A unique replicate identifier (sample_id) was generated for each cell by concatenating the disease group label (TB or HC) with the hashtag oligo identifier (HTO_maxID). Gene expression counts were aggregated at the sample level using the AggregateExpression function in Seurat (assay = “RNA”, layer = “counts”) to create a pseudobulk count matrix for each sample. Corresponding sample metadata containing group information was compiled and aligned to the count matrix to construct a DESeqDataSet object using the DESeq2 (v1.42.0). (Love et al., 2014) Genes with low counts (total count < 10 across all samples) were filtered out to improve statistical power. We set “ HC” as the reference group for comparison. DESeq2 was then used to normalize counts, estimate dispersion, fit models, and perform Wald tests for differential expression between the groups (TB and HC). Genes with an adjusted p-value < 0.05 and absolute log_2_ fold change > ±1.0 were considered significantly differentially expressed. Results were visualized using volcano plots generated by EnhancedVolcano (v1.20.0) and ggplot2 (v3.5.2), and key genes were labelled. The lists of significantly upregulated and downregulated genes were extracted, sorted, and exported for further interpretation.

### Mtb H37Rv culture

The Mtb H37Rv strain was revived from glycerol stocks stored at - 80°C. The stocks were inoculated in 7H9 media (BD Difco Middlebrook) supplemented with OADC (10%, Oleic acid, albumin, dextrose and catalase), glycerol (0.2%), and Tween-80 (0.05%) and passaged to secondary culture, which was grown to mid-log phase (OD ∼ 0.6). Secondary culture was harvested by centrifuging at 3,500 rcf for 10 minutes at room temperature, either for single-cell preparation or whole-cell lysate (WCL) preparation. Single-cell preparation was performed by passing the Mtb pellet through a 22-G needle and then through a 26-G needle, five times each. For Mtb whole cell lysate preparation, Mtb cultures were harvested at mid-log phase and subjected to heat inactivation at 90°C for 30 minutes with lysis buffer (8 mM EDTA, 50 µg/ml RNase A, 50 µg/ml DNase 1, 1× protease inhibitor in PBS) with 3 cycles of bead beating (Biospec Mini-Beadbeater-16) at 3650 oscillations/minute, with a 30- second on/off cycle. The lysed solution was centrifuged at 10,000 rcf at 4°C for 20 minutes, and the supernatant was collected and filtered with a 0.2 µM PES filter before leaving the BSL-3 laboratory. The protein concentration in the lysate was quantified using the BCA assay, and the quality was assessed by SDS-PAGE followed by silver staining. The lysate was stored at -20°C and used for stimulation experiments.

### Cell isolation, culture and infection for *ex vivo* experiments

Whole bone marrow (BM) cells were isolated from the femur and tibia of C57BL/6 mice (6-8 weeks old, male). BM cells were cultured in media containing DMEM, FBS (10%), and M-CSF (40 ng/ml) after RBC lysis. These bone marrow-derived macrophages (BMDMs) were harvested on day 7 using phosphate-buffered saline (PBS-EDTA, 0.5 mM) and seeded into 6-well plates (2 million cells per well). BMDMs were allowed to adhere to the plate by incubating overnight. BMDMs were infected with Mtb H37Rv at a 1:5 multiplicity of infection (MOI) by incubating at 37°C for 4 hours in the presence of 5% CO_2_. Infection was terminated by removing the media and washing the cells four times with PBS. At different time points, the cells were lysed with SDS (0.06%) and plated on 7H11 agar plates, and the CFUs were enumerated after 21 days post-incubation.

For the isotopomer experiment, BMDMs were infected with Mtb H37Rv at an MOI of 1:5, and after removing extracellular bacteria, 4 hours post-incubation, ^13^C_5_/^12^C_5_-Methionine-containing DMEM with dialysed FBS (10%) was added to the respective experimental and control wells. Metabolites were harvested at various time points by quenching the cells with chilled methanol. One set of BMDMs differentiated with L929-conditioning media was infected with Mtb H37Rv (MOI, 1:5), and proteins were harvested at 0, 4, and 24 hpi by lysing with RIPA buffer followed by acetone precipitation (4 volumes of ice-cold acetone). Extracted proteins were quantified using the BCA assay and stored at -80°C until future processing for proteomic profiling.

In a separate experiment, BMDMs, differentiated with L929 conditioning media (20%), and post-Mtb infection, were grown in DMEM media either supplemented with methionine (MS; 5×; 1 mM/150 mg/L), methionine restriction (MR; 0 mg/L) or control (30 mg/L). Cell viability was monitored at 0, 24 and 48 hpi using the trypan blue assay. Cell culture supernatants harvested at these time points from Mtb-infected BMDMs were filtered through a 0.22 µm PES filter and stored at -80°C for cytokine estimation. The previously described methods were followed for CFU enumeration. In a separate experiment, control and methionine-supplemented macrophages were treated with caspase inhibitor (VX765, 20 µM) and harvested at 24 hpi for western blot analysis and CFU assay.

### *In vivo* methionine supplementation experiment

C57BL/6 mice (6-8 weeks, ♂) were housed in the animal house facility at ICGEB, New Delhi. A group of mice received methionine in drinking water (∼2.5%, 3× extra of normal methionine in the diet) along with vehicle (2% sucrose). The control group received the vehicle in drinking water for two weeks. Feed and water intake were measured daily; body weight was measured twice a week; and random fasting (∼6 hours of fasting) was performed; and blood glucose was measured every week. An intraperitoneal glucose tolerance test was performed after two weeks of methionine supplementation. Briefly, mice were fasted for 6 hours, and glucose (1 g/kg body weight) was introduced intraperitoneally. Blood glucose was measured via the tail vein at different time intervals (0, 15, 30, 60, 90, 120 minutes) using a glucometer (Dr. Morepem BG-03 Gluco One). A group of mice from both groups were anaesthetized using 5% Isoflurane, and organs were harvested from the humanely euthanized mice sacrificed after two weeks of intervention. Serum was collected for metabolite and cytokine profiling by centrifuging clotted blood at 3,500 rcf for 20 minutes at 4°C. Images of gross tissue pathology were taken, and wet weights were recorded for the lungs, liver, and spleen. Single cell suspension was prepared from the harvested lungs, and spleen using collagenase (2 mg/ml) and DNase (1 mg/ml) in RPMI-1640 (2 ml) at 37°C for 30 minutes. The enzymatically treated tissues were passed through a strainer (70 µm) before centrifuging at 400 rcf for 10 minutes to collect the pellet. Ammonium chloride potassium (ACK) lysis buffer was added to the pellet, incubated for 2 minutes at room temperature and quenched by adding RPMI-1640. The cells were centrifuged, resuspended in RPMI-1640, and counted using a hemocytometer. From each sample, 10 million cells were incubated with an antibody cocktail (Live/dead fixable dye, CD45, CD11b, F4/80, SiglecF for macrophage panel and Live/dead fixable dye, CD3, CD4, CD8, CD44 for T-cell panel) for 45 minutes at 4°C. An aliquot of cells was incubated overnight with phorbol 12-myristate 13-acetate (5 ng, PMA) and ionomycin (0.75 µg) or Mtb-H37Rv whole cell lysate (10 µg/ml). After washing, these cells were incubated with brefeldin (5 µg) and monensin (1.3 µg) for 3 hours. The cells were washed with FACS buffer (1% FBS in PBS) and incubated with an antibody cocktail containing LIVE/DEAD and surface markers for 40 minutes on ice. After staining, these cells were fixed using paraformaldehyde (4%) and permeabilized using Permeabilization buffer (#00-8333-56, Invitrogen, USA). These cells were incubated with an antibody cocktail of intracellular cytokines: TNF-α, IL-17A and IFN-γ for 45 minutes, then washed with FACS buffer and finally resuspended in the FACS buffer. Flow cytometry data were acquired using BD LSR Fortessa X-20 or BD Symphony A1 and analysed using FCSExpress (DeNovo software, Version 6.0).

Post 2 weeks of methionine supplementation, the mice were transferred to the BSL-III tuberculosis aerosol challenge facility (BSL-3) at the host institute. These mice were aerosol challenged with a low dose of animal-passaged Mtb H37Rv (100-120 CFU, aerosol mode via Madison chamber). A subgroup of mice from the methionine-supplemented group was shifted to the control (MS to CT) post-Mtb infection. At the end of the study (21 dpi), the mice were humanely euthanised for tissue and biofluid collection. Harvested tissues (lungs, spleen, and liver) were processed using a homogenizer in sterile PBS (1 ml), and an aliquot (100 µl) of it was inoculated on 7H11 agar plates supplemented with OADC (10%), and BD MGIT™ - PANTA™ antibiotic mixture at appropriate dilution in sterile PBS and the colonies were counted post 3 weeks of incubation at 37°C in the incubator. Lung and spleen cells were processed for flow cytometry at the earlier time point. Serum was stored at -80°C and later on used for metabolite profiling and cytokine estimation. An aliquot of serum was used for Superoxide dismutase assay (E-BC-K019-M, ELabscience, China), Alanine aminotransferase (AST, #701640, Cayman Chemicals, USA) and Aspartate aminotransferase (ALT, #700260, Cayman Chemicals, USA) assays, performed according to the manufacturer’s protocols.

### Cytokine estimation

Immunoassay plates were coated with diluted capture antibody (100 µl, 200-fold diluted from a stock in assay buffer, i.e., 1% BSA in PBS) and incubated overnight at 4°C following the manufacturer’s protocol (BioLegend #432601). The plates were washed four times with washing buffer (0.02% Tween-20 in PBS; PBST) and blocked for non-specific binding with bovine serum albumin (BSA, 1%) for 1 hour at 25°C at 125 rpm. Standard (0 to 2,000 pg/ml) and samples were allowed to bind for 2 hours. The plates were washed and incubated with the detection antibody (100 µl, 200× diluted from stock in assay buffer) for an hour. After washing with PBST, plates were incubated with Avidin-HRP substrate (100 µl, diluted from 1,000× stock in assay buffer) for 30 minutes. The plates were developed by adding TMB substrate, and the reaction was stopped using 2N HCl. Absorbance was monitored at 450 nm and 570 nm using a multiplate reader, and the data were analysed in Microsoft Excel to identify group-specific differences, if any.

### Histopathology

Formalin-fixed lungs were paraffin-embedded, sectioned and stained with hematoxylin and eosin, followed by capturing images at 20× magnification. The slides were independently analysed by a histopathologist and calculated the total granuloma score, bronchitis, peribronchitis, perivasculitis, alveolitis, alveolar hyperplasia, capillary congestion, edema and fibrous tissue proliferation for lungs; lobular inflammation, portal inflammation, steatosis, infiltration, necrosis and hepatocyte degeneration for liver; cellular infiltration, congestion, lymphocyte apoptosis, necrosis, fibrosis, pigment and follicular hyperplasia for spleen.

### Metabolite extraction and analysis using LC-MS/MS

To the BMDMs harvested at different time points, chilled methanol (GC grade, 0.5 ml, 80%) was added to quench metabolic reactions and the samples were stored at -20°C until further processing. Culture media at these time points were collected and filtered through a 0.2 μm nylon filter before storage for extracellular metabolite profiling. For intracellular metabolite extraction, ribitol (2μl, 0.5 mg/ml) was added to each sample as a spike-in standard before incubation in a thermomixer (1,400 rpm, 4 °C, 30 minutes). The lysates were then centrifuged at 10,000 rcf for 10 minutes at 4 °C. The supernatant was collected and filtered through 0.2 μm filters. GC-grade chloroform (0.5 ml) was added to the filtered lysates, which were then vortexed briefly before centrifuging at 10,000 rcf for 10 minutes at 4 °C. The polar and non-polar phases were collected in fresh methanol-treated MCTs, and the interphase was left in the same tube. Quality control (QC) samples were prepared by pooling equal amounts of all samples (intercellular metabolites). The polar metabolites and interphase were dried in SpeedVac at 40 °C.

For metabolite isolation from lungs and liver, tissues (100 mg) were finely chopped and transferred to bead-beating tubes (2 ml) containing zirconium beads (2 mm, 250 mg). Chilled methanol (80%, 1 ml) was added to the tubes and ribitol (5 μl, 2 mg/ml) was added as a spike in standard. The tissues were subjected to bead beating using a Biospec Mini-Beadbeater-16 at 3650 oscillations/minute, with 30 seconds on/off cycles six times, with incubation on ice during the off cycle. After bead beating, the samples were incubated on ice for 30 minutes, followed by centrifugation at 10,000 rcf for 10 minutes at 4°C to collect the supernatant (800 μl). The extracted metabolites were filtered (0.2 μm, nylon membrane filter #726-2520), dried in a SpeedVac and stored at -20°C until further processing. An equal volume (120 μl) of the extracted metabolites from all the samples from a particular time point was pooled before drying to prepare a QC sample.

For serum metabolite extraction, chilled mass spectrometry grade methanol (80%) was added to serum (50 µl), which was then incubated on a thermomixer (900 rpm, 4°C) for 20 minutes. After centrifuging (10,000 rcf, 4°C for 10 minutes), the supernatant was collected, dried in a SpeedVac (Labconco® Refrigerated CentriVap® Benchtop Vacuum Concentrator, USA) and stored at -20°C until LC-MS/MS data acquisition. A quality control (QC) mixture was prepared before drying.

The metabolites were resuspended in buffer (5% acetonitrile, 5% internal standards, water), and metabolomics data were collected by injecting into the Thermo ScientificTM UHPLC system coupled with the Q Extractive Orbitrap mass spectrometer. Hypersil GOLD C18 Selectivity HPLC Column was used for the metabolite separation and set at 60 °C. The mobile phases consist of 1% formic acid (A) and 100% Acetonitrile (B). The LC gradient started with 5% ACN at a flow rate of 0.5 mL/minute, increased to 99% ACN for the next 20 min, then washed with 5% ACN for 5 min, for a total run time of 25 min. Ion source temperature ranged from 250-300 °C, and a spray voltage of 5500–4500 V. The data were acquired with MS1/MS2 of 150 to 2,000 m/z range, nitrogen as collision gas, and a resolution of 70,000 m/Δm. The MS data were acquired in positive and negative modes at ±80 eV in Orbitrap resolution with a mass range from 150 to 2,000 Da. From the Mtb-infected BMDMs, proteins were extracted and, after resuspending in TEAB (100 μl, 100 mM), quantified using the BCA assay for normalising metabolite peak intensity data.

### Metabolomics data analysis

The raw LC-MS/MS data files (n=51; 24 intracellular samples, 24 culture supernatant and 3 QCs) were processed through Compound Discoverer software (2.0, Thermo Fisher Scientific, USA). The raw MS/MS files were subjected to spectra selection and retention time alignment. The resulting spectra were used for compound identification via the Chemspider node, selecting databases (MMDB, HMDB, PubChem, DrugBank and LipidMaps). The mass tolerance for the compound detection node was set at <5 ppm with an S/N threshold of <3 (**Supplementary Figure S2A)**. The resulting matrix was down-selected by removing unknown hits, summing duplicate values for adducts of the same compounds, and manually removing contaminating features from solvents in Microsoft Excel (**Supplementary Figure S2B**). The final data matrix was normalised using an internal standard (dimetridazole) and protein amount, and was uploaded to MetaboAnalyst 6.0 for univariate and multivariate analysis. The plots generated in the MetaboAnalyst 6.0 were recreated using GraphPad Prism (v 8.4.2).

For isotopomer data analysis, a stable-isotope labelling workflow using an online database was employed. Raw MS/MS files for ^13^C samples were designated as labelled samples, and the ^12^C samples at the same time point were used as controls for identification. A blank run file was used for background correction, and standards were run to confirm peak identity and retention time. After retention time alignment and mass tolerance filtering, annotation with the ChemSpider node using the HMDB database, the Analyse labelled compounds node was used to obtain ^13^C labels for up to 25 exchanges. The results matrix was used to down-select these metabolites containing ^13^C carbons using a stable-isotope labelling layout. Relative exchange rates (%) for different isotopologues of the compounds of interest were plotted to compare carbon unit transfer from Methionine.

### Proteomics sample preparation and run using nanoLC-MS/MS

Equal amounts of harvested proteins (20 μg) from each experimental time point and conditions were resuspended in 100 mM TEAB buffer and reduced with 200 mM Tris(2-carboxyethyl)phosphine for 1 hour at 55 °C. Reduced proteins were alkylated with iodoacetamide (5 μl, 375 mM) at room temperature in the dark, and the reaction was dried in a SpeedVac (Labconco, USA) after centrifuging at 8,000 rcf at 4°C for 10 minutes. To the dried alkylated proteins, sequencing-grade trypsin (2.5 μg/100 μg of proteins, Promega, WI, USA) were added and incubated at 37°C overnight. Trypsin digestion was stopped by adding anhydrous acetonitrile, and the digested peptides were then dried in SpeedVac at 40 °C. The tryptic peptides were desalted using a ZipTip C_18_ column (Pierce, Thermo Fisher Scientific). Briefly, these columns were wetted with acetonitrile (100%) and equilibrated with formic acid (0.1%). Peptides were loaded onto the column by passing it 7-8 times and then washed twice with methanol (5%) containing formic acid (0.1%) to remove unbound impurities. Elution solution (50% acetonitrile in 0.1% formic acid) was used to elute the peptides. The eluted peptides were dried in SpeedVac and, at the time of the LC-MS/MS run, were resuspended in formic acid (0.1%, 15 μl).

Tryptic peptide fractions were separated on nano-LC Easy nLC 1200 (Thermo Fisher Scientific, Singapore) at a flow rate of 300 nl/minute on C18 precolumn (Acclaim PepMap 100, 75 μm × 2 cm, nanoViper, P/N 164,946, Thermo Fisher Scientific Incorporation, USA) followed by analytical column (Acclaim PepMap RSLC C18, 75 μm × 50 cm, 2 μm, 100 Å, P/N ES803) using a gradient of solvent B (95% acetonitrile in 0.1% formic acid) from 5% - 10% for 0 to 5 minutes followed by 35% for 5 to 95 minutes and finally, 95% for 95 to 130 minutes and solvent A (0.1% formic acid). The eluted peptides were injected into the mass spectrometer (Orbitrap Fusion Lumos Tribrid Mass Spectrometer), and MS1 data were acquired in positive-ion mode at 120,000 Orbitrap resolution, with a mass range of 375 to 2,000 Da. Precursor ions were allowed to fragment using a higher-energy C-trap dissociation (HCD) in an ion trap (IT) detector with a collision energy of 38% in a data-dependent MSn Scan acquisition at a resolution of 50,000. Precursor ions with +2 to +6 charge and monoisotopic ions were selected. Parent ions, once fragmented, were excluded for 60 s with an exclusion mass width of ± 10 ppm.

All tandem mass spectra data were analysed using Proteome Discoverer software (version 2.3, Thermo Fisher Scientific). The raw MS/MS data files were searched against the *Mus musculus* UniProt proteome database (ID: UP000000589; accessed on 28/01/2024; 54,739 proteins) and contaminant protein database (PD_Contaminants_2015_5.fasta, provided by the manufacturer). A maximum of two trypsin cleavages was allowed with a precursor mass tolerance of 10 ppm and fragment mass tolerance of 0.8 Da. Carbamidomethylation (+57.021 Da, Cys) at the C termini was selected as a static modification. Oxidation +15.995 Da (Met) and acetylation +42.011 Da at the N-terminus were selected as dynamic modifications, and proteins were identified based on at least two peptides at an FDR < 0.05. Mascot and SequestHT were used as analysis tools to assign identities to spectra via precursor-ion quantification of proteins. Proteins that met the criteria, qualifying the parameters, like log_2_ fold change ≥ ±1.0; -log_10_ adjusted p-value ≥ 1.3 (p-value<0.05) between study groups, were selected as significantly deregulated molecules.

### Elements quantification using an Inductively coupled plasma-mass spectrometer

To the harvested serum (25 µl), HNO_3_ (70%, 75 µl) was added and transferred to sample vials (MG5, Anton Paar, USA), and then H_2_O_2_ (30%, 25 µl) was added. The sealed sample vials were subjected to microwave (Anton Paar, USA) digestion using a ramp to 250W for 15 min, with a 5 min hold at 250W. The initial power was set at a ramp of 10 minutes and 150W with a max temperature of 140°C, followed by a 15-minute hold. Then, a second ramp of 15 minutes from 150 to 250W was used, followed by a final hold to bring the temperature to 55°C at the highest fan speed. After diluting the samples with trace metal-free water (Honeywell International Inc.), element levels (^24^Mg, ^44^Ca, ^57^Fe, ^63^Cu, ^66^Zn, ^77^Se) were monitored in the digested samples using an Inductively Coupled Plasma- Mass Spectrometer (iCAPTM TQ ICP-MS, Thermo Fisher Scientific, USA). Thermo Scientific Qtegra^TM^ Intelligent Scientific Data Solution (ISDS) software was used to operate and control the instrument. Briefly, the torch was warmed up for 30 minutes in single-quad Kinetic energy discrimination (SQ-KED) mode with helium to remove polyatomic nuclear interference, and then autotuned in normal mode, followed by advanced KED mode. Then, the multi-element standard (#92091, Sigma Aldrich, USA) at different concentrations was run under the same conditions to prepare the standard plot. A sample blank was run before the run. Each element was quantified using a standard plot, and group-specific details were compared. Data analysis was done using Qtegra software (Thermo Fisher Scientific, USA).

### Chromatin immunoprecipitation quantitative PCR

BMDMs (1×10^7^) were seeded in 90 mm dishes and cultured in DMEM supplemented with 5× Methionine (MS) or in a non-supplemented group as Control (CT). BMDMs were infected with the Mtb H37Rv strain at an MOI of 1:5. At 24 hpi, cells were crosslinked with formaldehyde (1% for 10 minutes at room temperature). Cross-linking was terminated with Glycine solution (1×), then cells were lysed using Buffer A and Buffer B provided with the ChIP kit (SimpleChIP Plus Enzymatic Chromatin IP Kit, Cell Signalling Technologies, CST#9005). Chromatin solutions were sonicated to yield DNA fragments of 150-900 bp using a Sonics Vibra cell sonicator (VCX-130 sonicator, Sonics and Materials Inc., USA). Sonicated chromatin was incubated with monoclonal mouse H3K4me3 and H3K36me3 antibodies (Abcam, #ab8580 and #ab9050) overnight at 4°C. As a control, sonicated chromatin was incubated with IgG antibodies, and 2% of the chromatin was used as an input control. Protein G magnetic beads were used to enrich the Ab-Histone complexes. Bound beads were washed thrice with low-salt wash buffer and once with high-salt wash buffer. Immuno-complexes were eluted with the elution buffer. Cross-links were reversed in the final volume of 200 µl using NaCl and proteinase K treatment for 2 hours at 65°C. DNA was purified using the DNA purification Spin columns provided with the kit. Enrichment was measured by qPCR of DNA immunoprecipitated with anti-H3K4me3, anti-H3K36me3 and IgG using the primers indicated (**Table 1**). Fold enrichment for the relative gene expression was calculated using the comparative threshold cycle method.

**Table 1:**
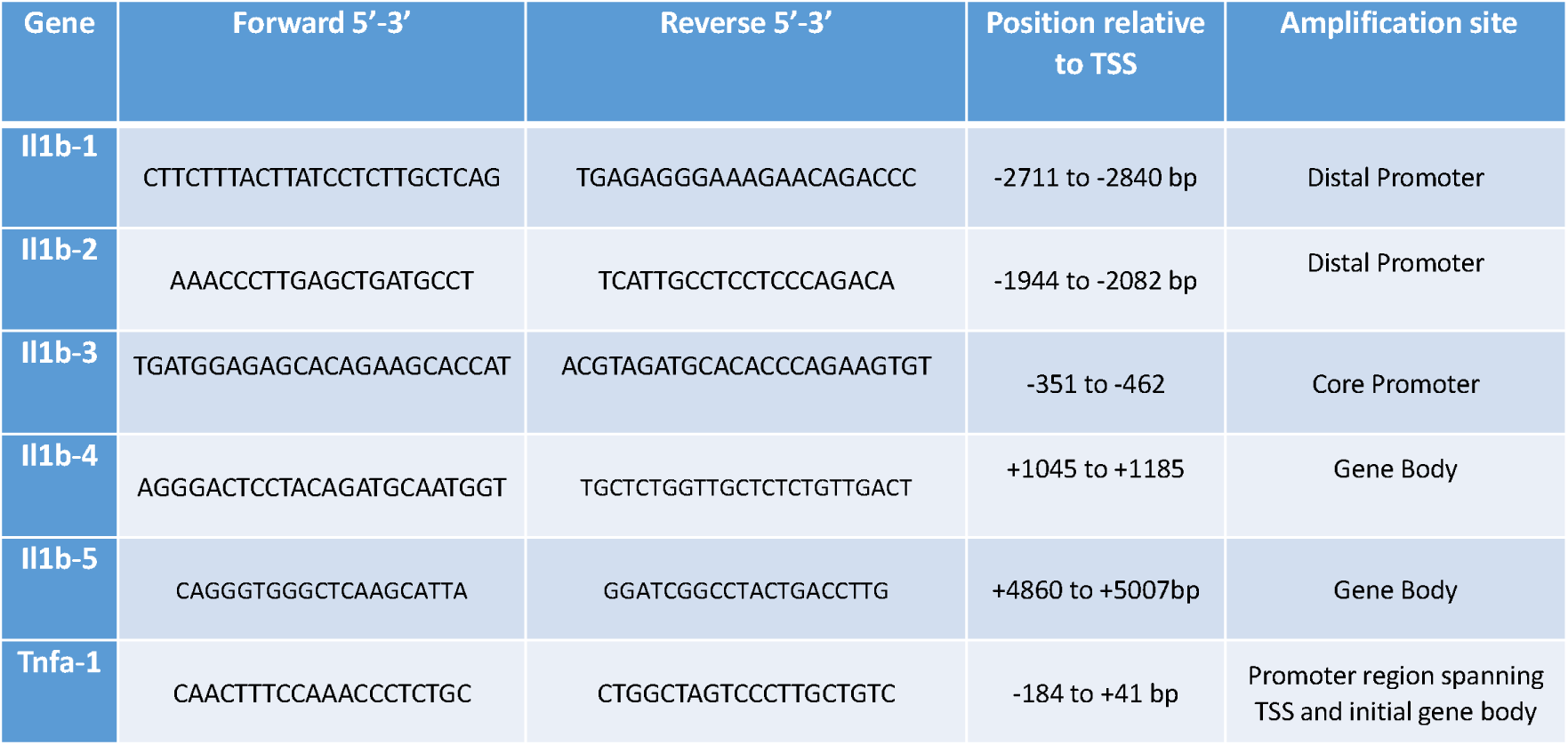
Primers used for ChIP qPCR for the Il-1b and Tnf-a genes.

### Statistical analysis

Raw LC-MS/MS files were analysed using Compound Discoverer, and MetaboAnalyst 5.0 was used for multivariate and metabolite set enrichment analysis (MSEA) of metabolomics data. Metabolite and protein abundances between study groups were compared using the Mann-Whitney t-test in GraphPad Prism. Student’s t-test, Welch’s t-test and one-way ANOVA were used for comparing CFU, cell viability and cytokine data and p-value<0.05 or log_2_FC ±1.0 and log_10_p-value>1.3 are considered for a significant difference. Linear regression/4-PL plots were used for calculating unknowns for ELISA and targeted metabolomics after standard curve generation. For elemental analysis, concentrations were determined from respective standard curves, and group-specific differences were compared using One-way ANOVA; Tukey’s multiple-comparison test was used to calculate p-values.

## Results

### scRNAseq analysis of lung immune cells from Mtb-infected mice showed functional heterogeneity in the macrophage population

Macrophages, upon mycobacterial infection, exhibit perturbed tryptophan, glutamine and arginine metabolism, with limited literature on its impact on methionine metabolism. In our earlier report, we analysed the scRNAseq data of Mtb-infected C57BL/6 mice and control groups and identified three clusters of lung macrophages at 21 dpi (**Figure 1A** and **1B**, Chaudhary et al., 2025). The two clusters of alveolar macrophages (AM1 and AM2) showed distinct phenotypes: AM2 expanded from monocytic precursors, whereas AM1 represented the original tissue-resident population, which is widely shown to harbour Mtb (**Figure 1C** and **1D**). The Mtb-infected mice’s lungs had higher numbers of AM2 and non-AMs with minor change in the AM1 numbers, where the expanding macrophages show better pro-inflammatory phenotype with higher *Il-1b, Hif-1a* and *Nos2* expression (**Figure 1E**). The TB group macrophages showed increased *Il-1b* transcript levels with minimal expression in the AM2 population, highlighting it as a permissive niche for Mtb (**Figure 1F** and **1G**).

**Figure 1:**
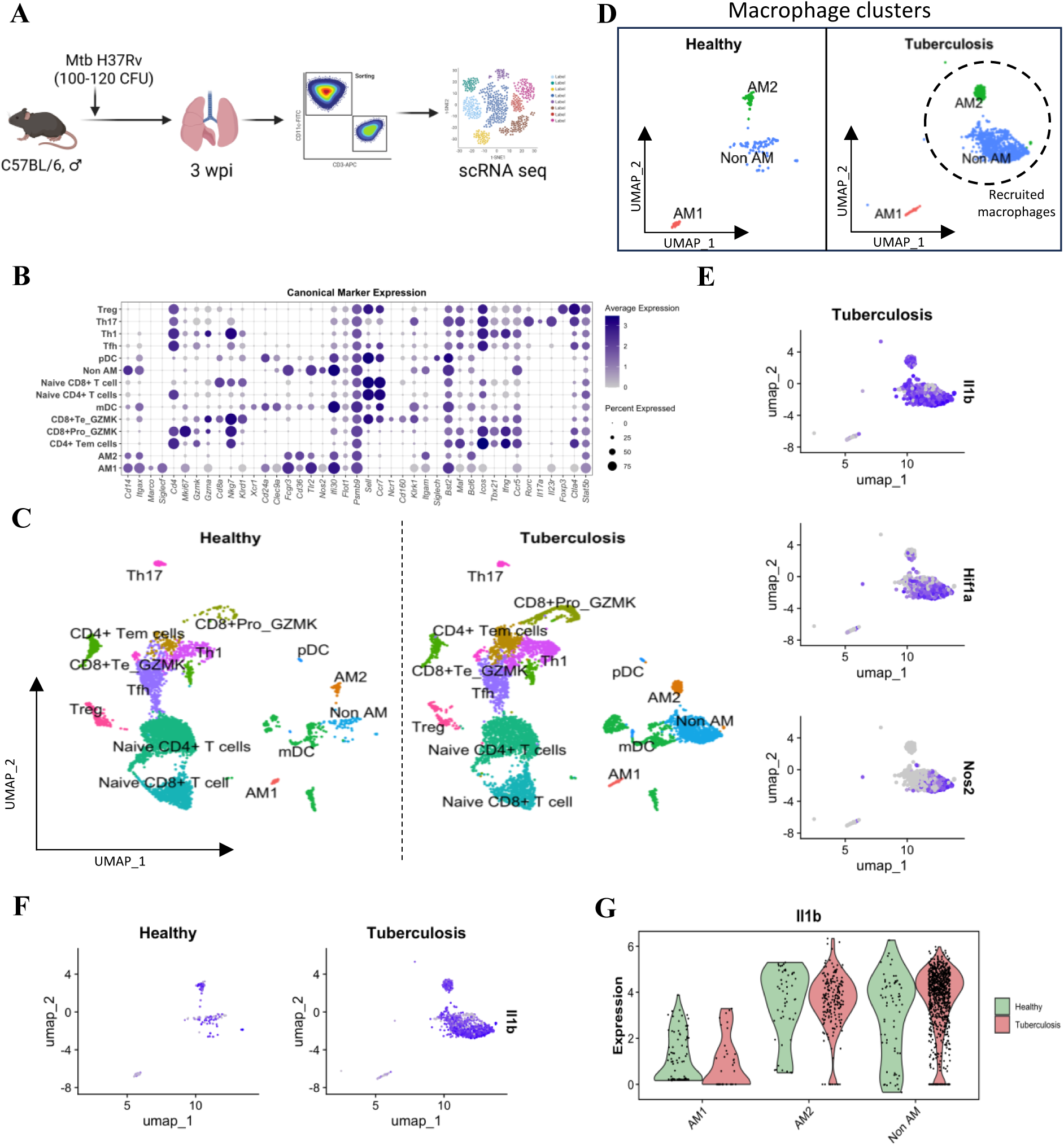
*Mycobacterium tuberculosis* H37Rv-infected C57BL/6 mice showed increased expansion of IL-1β-producing monocyte-derived macrophages. **A.** Methodology adopted for the scRNAseq experiment. **B.** Dot plot indicating expression of key genes used to classify each cell subtype. **C.** UMAP plots showing increased frequency of alveolar and non-alveolar macrophages in the Mtb-infected group. **D.** AM2 and Non-AMs show expansion in the lungs of the TB group. **E.** UMAP plots showing expression of *Il-1b, Hif1a* and *Nos2* genes in macrophage clusters in the TB group. **F.** UMAP plot showing *Il-1b* expression in macrophage clusters of TB and the healthy group. **G.** Violin plot showing *Il-1b* gene expression in various macrophage subclusters. TB: Tuberculosis, Mtb: *Mycobacterium tuberculosis*, scRNAseq: single cell RNA sequencing, AM: Alveolar macrophage.

To monitor functional heterogeneity among the three macrophage clusters in the TB group, if any, at the transcriptomic level, dysregulated transcripts (DEGs; log_2_FC >±1.0) were identified. AM1 cluster showed alterations in IL-1β response, one-carbon and glutathione metabolism related transcripts (IL-1 response: *Il-1b, Gsdmd,* One-carbon metabolism: *Mthfr, Mthfd, Dnmt3a, Msrb2, Phgdh,* Glutathione metabolism: *Gstt1, Gsto1, Gstm1)* (**Supplementary Figures S1A, S1B** and **S1C**). Our observation of compromised inactive glycine/serine biosynthesis in alveolar macrophages of Mtb-infected mice corroborates earlier reports. (Kurita et al., 2021)(Gan et al., 2021) We also observed significantly higher expression of Slc7a5, the neutral amino acid transporter responsible for extracellular methionine uptake in macrophages, and of *Msrb* and *Msra,* which are responsible for salvaging methionine from oxidised proteins in alveolar macrophages, highlighting the decreased availability of methionine. (Yoon et al., 2018)(Pandit et al., 2023) IL-1 responses, especially IL-1β signalling, are critical for host resistance and antimicrobial activity in macrophages via inflammasome activation and triggering TNF-α and IFN-γ production in the early phase of infection against Mtb. (Jayaraman et al., 2013)(Yamada et al., 2000) These alveolar macrophages are among the first immune cells to encounter Mtb and are permissive for Mtb growth, providing a niche for the bacilli to grow. (Cohen et al., 2018) We observed a decrease in the active IL-1 signalling from the AM1 cluster towards CD4+ Tem and Th17 cells in the TB group (**Supplementary Figures S1D and S1E**).

However, non-alveolar macrophages are considered restrictive to Mtb growth and play an important role in controlling Mtb infection. (Pisu et al., 2021)(Huang et al., 2018) Along with higher *Il-1b* expression, we observed a prominent infection-induced metabolic shift leading to an increase in glycolysis in the non-AMs, which is responsible for the pro-inflammatory phenotype in macrophages (**Supplementary Figures S1F** and **S1G**). These findings highlight that downregulation of methionine availability and serine synthesis may be leading to altered IL-1 responses in AM1 cells, thereby promoting bacterial survival in the alveolar macrophages. These findings suggest compromised infection-induced metabolic remodelling and perturbed one-carbon metabolism, which impact the IL-1 response and mycobacterial growth in permissive alveolar macrophages.

### Mtb-infected bone marrow-derived macrophages (BMDM) show perturbed methionine metabolism

For a molecular-level understanding of the Mtb infection and its impact on the one-carbon and the amino acid metabolism in the early phases of infection, M-CSF-differentiated macrophages from the bone marrow precursors of C57BL/6 mice were infected with Mtb H37Rv strain at an MOI of 1:5. At different time points, these Mtb-infected macrophages were subjected to CFU assay, and extracted metabolites were subjected to global metabolite profiling using LC-MS/MS (**Figure 2A, Supplementary Figure S2A a**nd **S2B)**. Mtb-infected macrophages showed similar viability till 24 hours post-infection (hpi), and the mycobacterial burden showed a significant increase at 24 hpi (**Figure 2B**). Global metabolite analysis of these Mtb-infected BMDMs and controls identified 371 putatively annotated metabolic features, and the PCA plot showed a tight cluster of all the QC samples, indicating minor contribution from the adopted experimental procedures (**Figure 2C**). At early time points (i.e. 0, 4 hpi), the metabolic profiles of the Mtb-infected macrophages showed overlapping clusters, whereas those clustered away at 24 hpi, highlighting a major metabolic shift (**Supplementary Figure S3A**). At 0 hpi, the metabolic profile of the Mtb-infected BMDMs clustered away from the uninfected controls, with a set of 172 deregulated (log_2_ fold change ≥ ±1.0; p-value < 0.05) metabolites (164/6: higher/lower), showing Mtb infection-associated metabolic variations (**Supplementary Figure S3B**). Pathway enrichment analysis of the identified deregulated metabolites showed a perturbed purine, glutathione, glycine and serine metabolism (**Supplementary Figure S3C**). Higher levels of adenine, spermidine and guanine were observed in Mtb-infected macrophages compared with uninfected controls. So, Mtb-infection had a significant impact on macrophage amino acid metabolism.

**Figure 2:**
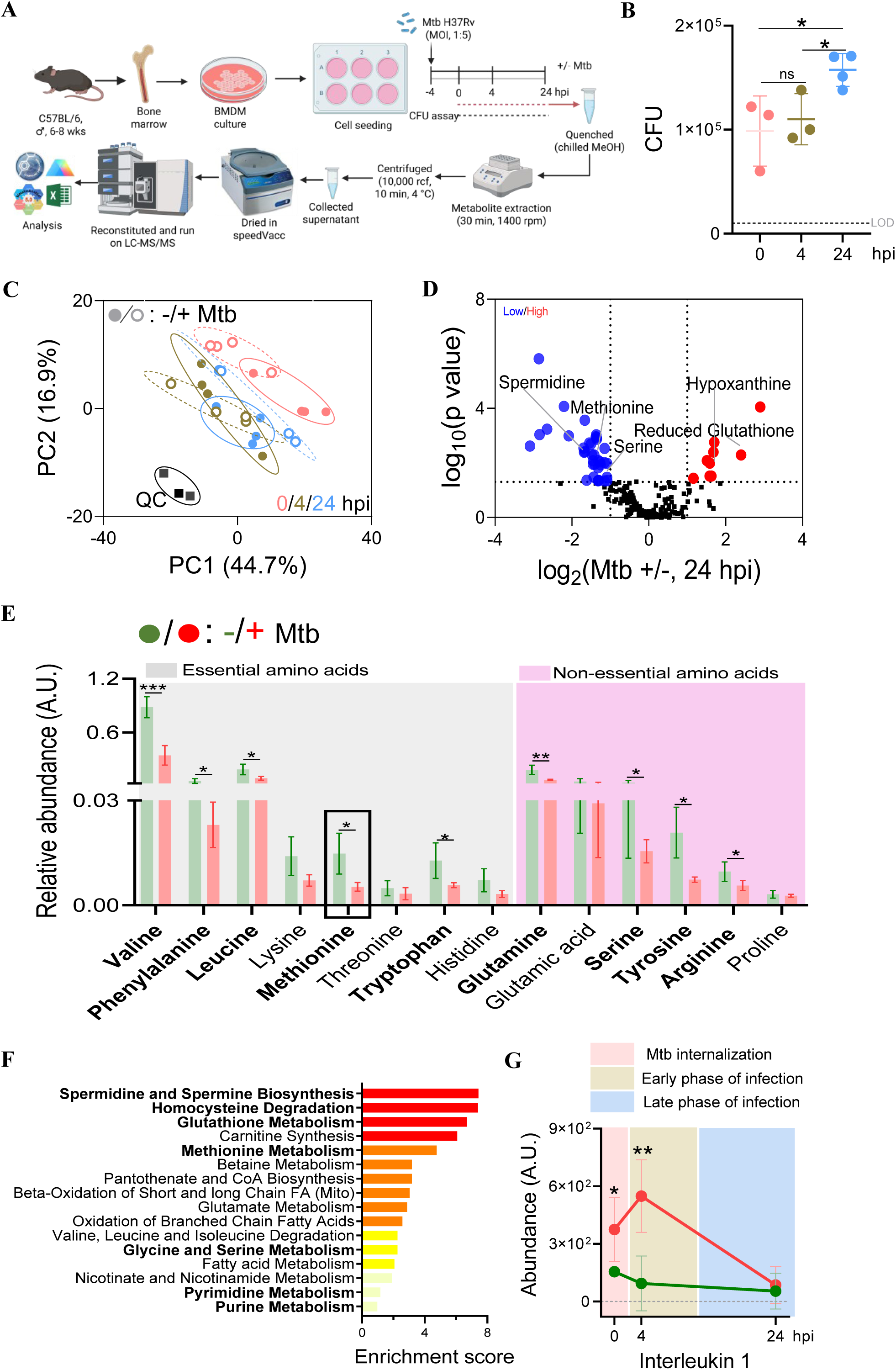
Mtb H37Rv-infection in bone marrow-derived macrophages (BMDMs) influences methionine and serine metabolism and IL-1 response. **A.** Schematic representation of the adopted experimental protocol. **B.** Intracellular Mtb load in BMDMs as assessed via CFU assay with dashed line showing the limit of detection (LOD). **C.** Principal component analysis (PCA) plot showed metabolome changes in the BMDMs across time points upon Mtb H37Rv infection and uninfected (+/-) and quality controls (QC). **D.** Volcano plot showing the deregulated metabolites of Mtb-infected BMDMs vs uninfected control at 24 hpi. **E.** Relative abundances of essential and non-essential amino acids in Mtb-infected macrophages compared with uninfected controls. **F.** Pathway enrichment analysis plot presenting significantly altered metabolic pathways at 24 hpi. **G.** IL-1β levels in Mtb-infected macrophages and uninfected controls at 0, 4 and 24 hpi. One-way ANOVA with Tukey’s multiple comparison is used to compare CFU data, n=4/group/time point, ns: non-significant at 95% confidence, *: p value< 0.05. +Mtb: Mtb-infected, -Mtb: Uninfected controls, hpi: hours post infection.

### BMDMs switch from pro-inflammatory phenotype-associated metabolic changes in the early phase to anti-inflammatory changes in the late phase of Mtb infection

In the early phase of Mtb infection in macrophages is associated with a metabolic shift from oxidative phosphorylation to HIF-α-mediated glycolysis and an NF-κB-mediated pro-inflammatory response. At the early time point (i.e. 4 hpi) of Mtb infection, the metabolic profile of the Mtb-infected macrophages showed overlap with the uninfected controls with a set of 22 deregulated (18/4: higher/lower) metabolites (**Supplementary Figure S3B**). Significantly higher levels of spermine and spermidine, which act as pro-inflammatory metabolites, and decreased adenine levels, which are reported to be anti-inflammatory, were observed. Polyamines exhibit immunomodulatory properties, including engulfment of apoptotic cells upon infection. (McCubbrey et al., 2022) These deregulated metabolites in Mtb-infected macrophages showed enrichment of methionine, purine and polyamine metabolism (**Supplementary Figure S3C**).

However, at later time points, i.e. 24 hpi, the metabolic profiles of Mtb-infected macrophages and controls showed two non-overlapping clusters (**Supplementary Figure S3A**). A set of 33 deregulated (log_2_fold change ≥ ±1.0; p-value <0.05; 5/28: higher/lower) metabolites was identified in Mtb-infected cells compared to uninfected controls (**Figure 2D**). At 24 hpi, the Mtb-infected macrophages had significantly lower levels of methionine, serine and spermidine and higher levels of hypoxanthine, glutathione, and adenine (**Figure 2E**). Methionine leads to cysteine production via the transsulfuration pathway, and both serine and cysteine then contribute to glutathione production, which helps to clear bacteria. (Bhatia et al., 2020) Methionine levels are reported to be lower in the serum of active TB patients than in those of latent TB and healthy controls. (Cho et al., 2020) Pathway enrichment analysis of these deregulated metabolites revealed perturbations in polyamine metabolism and in nodes of one-carbon metabolism, including glycine, serine, methionine, polyamine, and homocysteine metabolism (**Figure 2F**). Downregulation of tyrosine, valine, glutamine and tryptophan metabolism was observed in the Mtb-infected group (**Figure 2F**), corroborating earlier published reports. (Borah Slater et al., 2023)(Jiang et al., 2022)(Borah et al., 2019)(Xiao et al., 2022)(Jiang & Shi, 2021) As macrophages provide a favourable niche for Mtb and mount a poor immune response against Mtb, we monitored IL-1 levels, which were significantly higher only in the early phase of infection. Caspase 1, which converts pro-IL-1β, a major pro-inflammatory cytokine, to IL-1β, and its levels were significantly higher in the early phase of Mtb-infected macrophages (**Figure 2G, Supplementary Figures S4A, S4B** and **S4C)**. At the early phase of infection (4 hpi), serine, methionine, and polyamine metabolism were transiently increased, and at the later phase (24 hpi), serine and methionine levels decreased significantly. However, metabolites of the nucleotide salvage pathway, including adenine, guanine and hypoxanthine, showed an inverse relationship with the serine and methionine levels **(Figure 3**). This observation demonstrates that, by 24 hpi, a flux of metabolites from one-carbon metabolism is diverted towards nucleotide salvage and glutathione production in Mtb-infected macrophages. Recent reports have demonstrated that nucleotides, primarily adenine, act as anti-inflammatory cytokines and dampen the immune response in macrophages. (Harber et al., 2024) So, Mtb-infected BMDMs showed a time- dependent increase in mycobacterial load, as observed in the CFU data, with the highest load at 24 hpi, and significantly impacted methionine and homocysteine metabolism.

**Figure 3:**
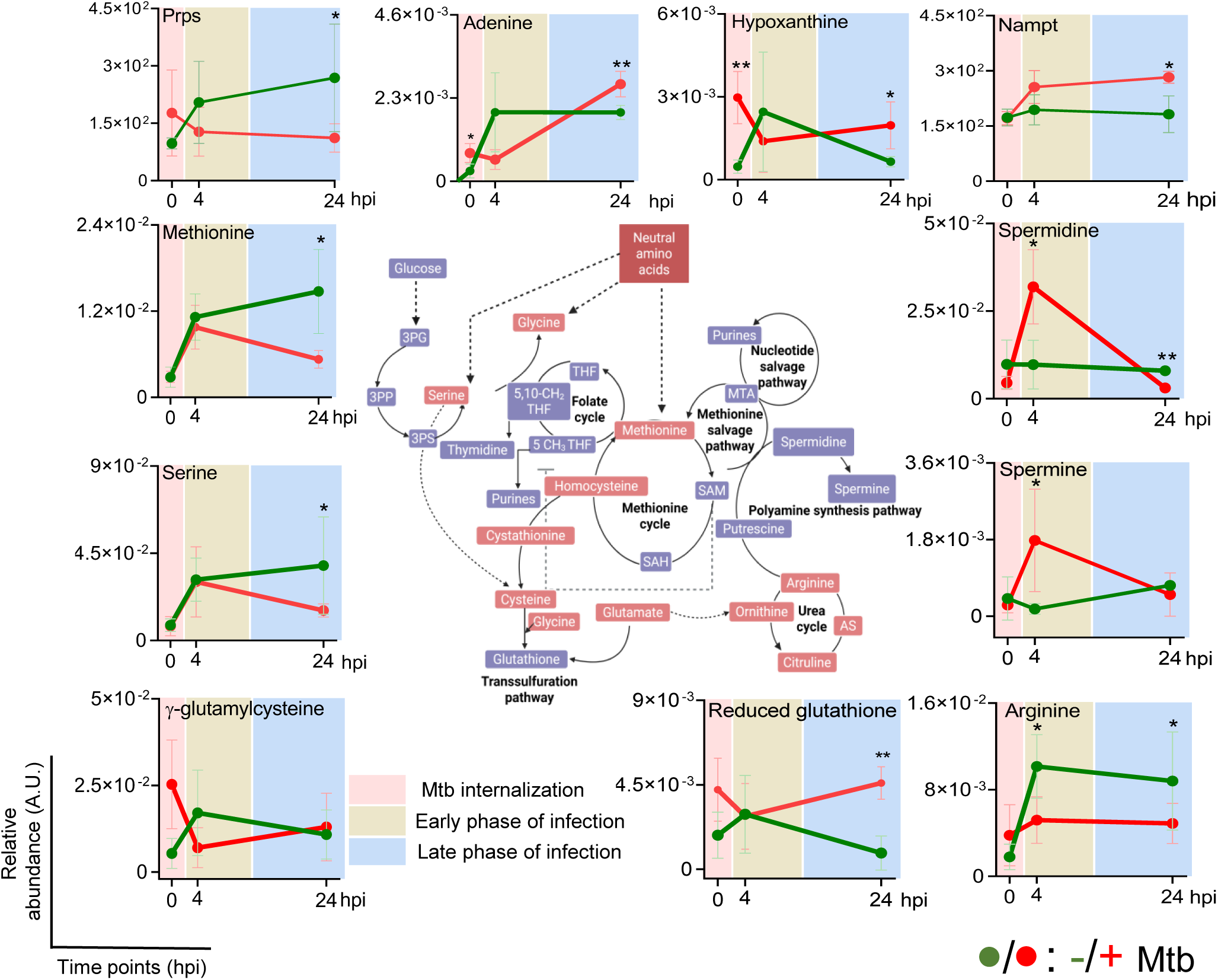
Kinetics of the metabolites related to the methionine-serine metabolic pathways and enzymes for nucleotide synthesis in the Mtb H37Rv-infected bone marrow-derived macrophages (BMDMs) and controls. BMDMs: Bone marrow-derived macrophages, Mtb: *Mycobacterium tuberculosis,* Prps: Phosphoribosyl pyrophosphate synthetase, Nampt: Nicotinamide phosphoribosyltransferase, n=4/group/time point, hpi: hours post infection; A.U.: arbitrary unit. *: p value< 0.05, **: p value<0.01.

### Mtb-infected macrophages show increased nucleotide salvage

We monitored the expression of proteins involved in one-carbon metabolism using proteomics data from these samples and time points. The enzymes involved in methionine and folate metabolism show similar abundance between the Mtb-infected and uninfected controls. Interestingly, enzymes involved in pathways diverting the one-carbon pool away from S-adenosyl methionine (SAM) showed significant alterations. Mtb-infected macrophages had significantly low Spermine synthase levels, which further reduced at later phases of infection (**Supplementary Figure S4D)**. Glutathione reductase levels were higher in the Mtb-infected macrophages, and arginase levels were significantly lower in the Mtb-infected group **(Supplementary Figures S4E** and **S4F**). Metabolite profiling showed elevated nucleotide levels, suggesting that carbon units from the methionine cycle are diverted to nucleotide salvage. Hence, we compared the expression of the rate-limiting enzymes involved in nucleotide salvage and de novo synthesis. Significantly higher levels of Nicotinamide phosphoribosyltransferase with similar expression of S-methyl-5’-thioadenosine phosphorylase were observed (**Figure 3**). Lower Phosphoribosyl pyrophosphate synthase levels in Mtb-infected macrophages might contribute to lowered de novo nucleotide synthesis (**Figure 3)**. Hence, increased nucleotide salvage from methionine metabolism could be correlated with a dampened IL-1 response and, thereby, a higher mycobacterial burden.

### Carbon units from Methionine metabolism get diverted towards polyamine metabolism, supporting nucleotide salvage

Methionine, being an essential amino acid, is sourced from extracellular sources in mammals, and the carbon units can be recycled via methionine salvage or by transfer of a methyl group from mTHF to homocysteine. Carbon units from methionine either lead to SAM production with the help of ATP or get diverted towards nucleotide salvage via the polyamine synthesis node (**Figure 4A**). To assess the fate of carbon units from methionine, Mtb-infected macrophages were cultured in media containing ^13^C_5_ Methionine. Intracellular levels of labelled methionine (M+5), spermine (M+3, M+6) and spermidine (M+3) were monitored using LC-MS/MS. Extracellular methionine was rapidly taken up by macrophages with 100% internalisation at 4 hours (**Figure 4B**). The relative abundance of total intracellular methionine levels corroborated with earlier experiments, showing decreased levels in the Mtb-infected group at the later phase of infection (12, 24 hpi; **Figure 4B**). The intracellular methionine pool showed similar levels at 4 hpi, a marginal increase at 12 hpi and a significant reduction at 24 hpi in the Mtb-infected macrophages, indicating diversion of carbon units or contribution of ^12^C Methionine either from protein degradation from the dead cells or bacteria engulfed by the macrophages (since ^12^C Methionine was seen in the extracellular counterpart) at the later phase of infection (**Figure 4C)**. Incorporation of ^13^C was higher in spermine (M+3) from the Mtb-infected group at 12 and 24 hpi (**Figure 4D**). However, ^13^C incorporation in spermidine was similar in both groups. We did not observe any ^13^C incorporation into nucleotides or glutathione since the carbon from methionine does not directly enter these metabolites. These data indicate that carbon units from methionine are diverted toward polyamine synthesis, which supports nucleotide salvage and thereby reduces the carbon available for methylation reactions via SAM.

**Figure 4:**
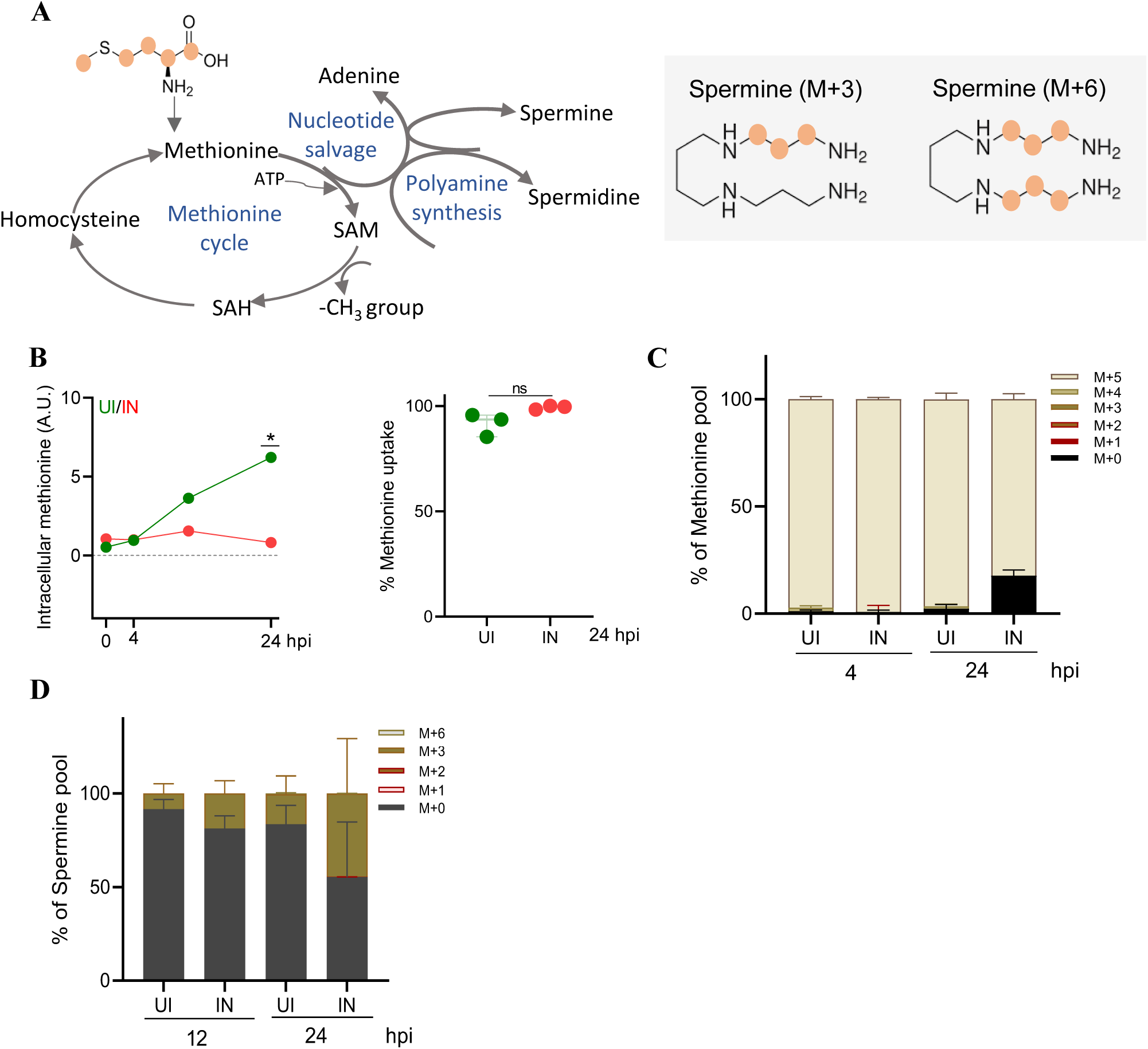
Mtb H37Rv-infected macrophages show diversion of carbon units from SAM towards polyamine synthesis, contributing to the nucleotide salvage pathway. **A.** Methionine cycle and its diversion node. **B.** Intracellular methionine pool and methionine consumption from media in Mtb H37Rv-infected and uninfected macrophages. The first plot shows the trend in the intracellular methionine pool in Mtb-infected and uninfected macrophages over time, and the second plot shows Methionine uptake calculated from the culture media at 24-hour intervals. **C.** Mass isotopomer distribution (MID) of intracellular methionine pool in Mtb-infected and uninfected macrophages cultured in ^13^C_5_-Met containing media for 4 and 24 hours. **E** and **D.** Mass isotopomer distribution (MID) of intracellular spermine pool in Mtb-infected and uninfected macrophages cultured in ^13^C_5_-Met containing media at 12 and 24 hpi. IN: Mtb H37Rv-infected macrophages, UI: Uninfected control macrophages, Mtb: *Mycobacterium tuberculosis*, Met: Methionine, hpi: hours post-infection, *: p-value < 0.05 at 95% confidence, MID data represents Mean ± SEM for biological replicates.

### Methionine supplementation increases the pro-inflammatory response in Mtb-infected macrophages

Essential amino acids like methionine primarily contribute to one-carbon metabolism, which is known to critically control the production of pro-inflammatory cytokines like IL-6, TNF-α and IL-1β. Other amino acids, like serine from either exogenous source or from the glycine cleavage pathway, also contribute to a minor proportion of one-carbon units for one-carbon metabolism and pro-inflammatory cytokine production. However, exogenous methionine supplementation (MS) or restriction (MR) might directly impact the pro-inflammatory response in Mtb-infected macrophages. Dietary methionine restriction has been reported to suppress inflammation in various immune cells, such as macrophages and T cells. (Roy et al., 2020) Methionine supplementation, however, is reported to improve alveolar macrophage function in patients with alveolar proteinosis and to induce M1 polarisation, thereby improving the immune response against Mtb. (Franceschi et al., 2020)(Santos et al., 2017)(W. W. Zhang et al., 2024) Importantly, supplementing methionine will not be beneficial for Mtb directly since Mtb synthesises its own methionine and S-adenosyl methionine and does not rely on host sources for an autarkic lifestyle. (Berney et al., 2015) Hence, we monitored the impact of methionine levels on the bacterial clearance and Mtb H37Rv-infected BMDMs cultured in media supplemented with methionine (1.0 mM, 5× concentrations) showed significantly higher viability as well as mycobacterial clearance compared to the methionine-restricted (0 mM) or control (0.2 mM) groups (**Figures 5A**, **5B** and **5C**). As reported earlier, methionine restriction compromised the BMDMs’ viability **(Figure 5B)**. Interestingly, methionine supplementation or restriction had a limited impact on cell proliferation up to 48 hours of culture, so the observed changes in Mtb CFU were not due to compromised cell viability (**Supplementary Figure 4G**). So, it seems, irrespective of sex, methionine supplementation has a beneficial antimycobacterial effect.

**Figure 5:**
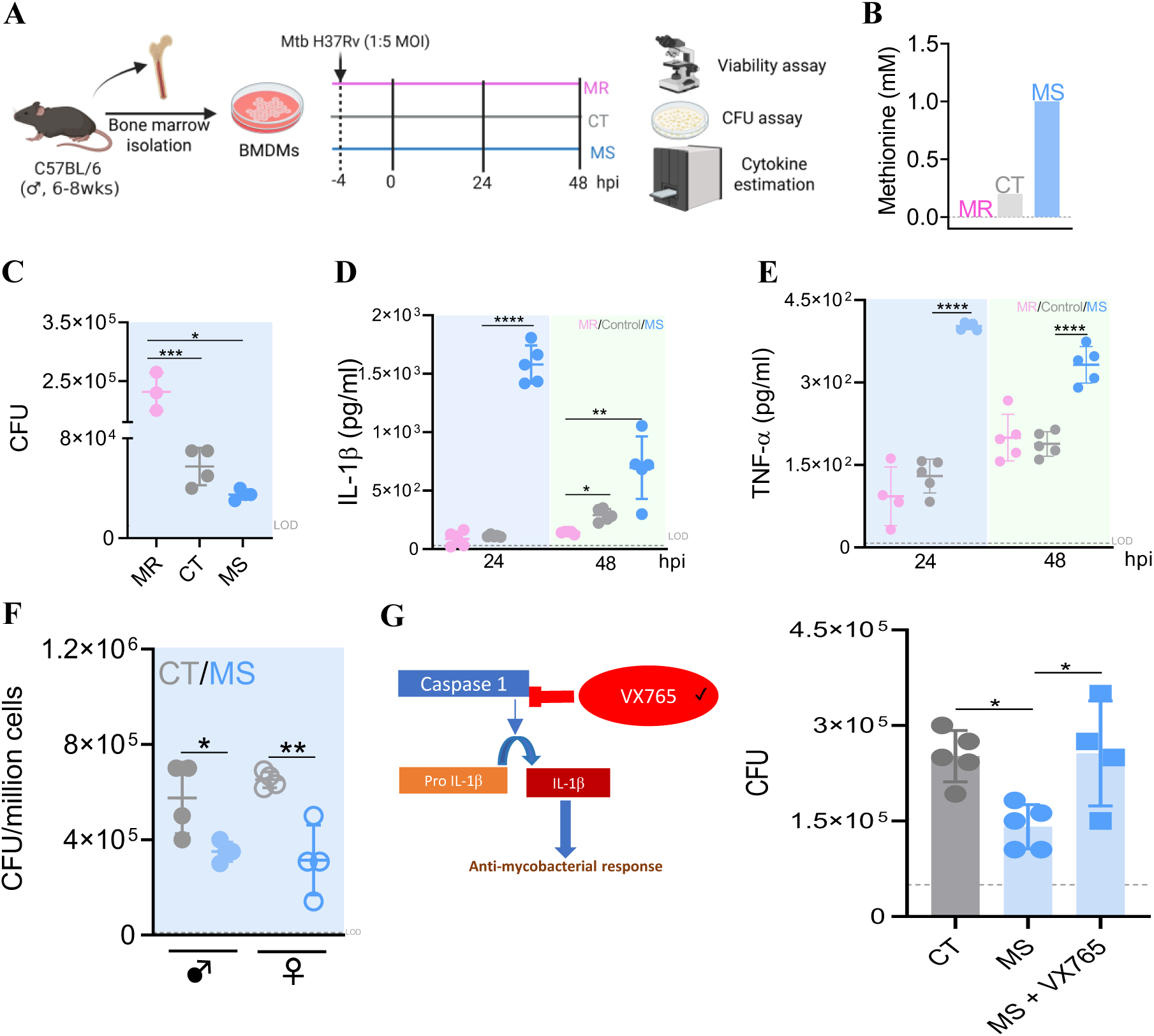
Methionine availability affects inflammatory profile and mycobacterial clearance in the bone marrow-derived macrophages (BMDMs). **A.** Schematic showing the experimental strategy adopted in this study. **B.** Methionine amount in culture media in different groups. **C.** Mycobacterial load in the macrophages, restricted or supplemented with methionine (MR/MS) and normal media (control) at 24 hpi. IL-1β (**D)** and TNF-*α* (E) secreted by Mtb H37Rv-infected BMDMs grown in restricted or supplemented with methionine and control at 24 and 48 hpi, respectively. **F.** Mycobacterial load in methionine-supplemented and control macrophages from male and female mice. **G.** Mycobacterial burden in macrophages treated with caspase 1 inhibitor (VX765). One-way ANOVA with Tukey’s multiple-comparison test is used to compare CFU and cytokine data. BMDMs: Bone marrow-derived macrophages; *: p-value<0.05; **: p-value<0.01; ***: p-value<0.001; ****: p value<0.0001 at 95% confidence. The dashed line represents the limit of detection (LOD).

The methionine-supplemented (MS) BMDMs, upon Mtb infection, released higher IL-1β at 24 and 48 hpi (**Figure 5D**). Interestingly, IL-1β production in the methionine-supplemented group decreased from 24 to 48 hpi, indicating that host cells exhibit an acute, controlled, but not sustained, pro-inflammatory profile. The methionine-supplemented group also showed consistently higher TNF-α production at 24 and 48 hpi as compared to the Mtb-infected control group (**Figure 5E**). The reduction in bacterial burden in methionine-supplemented macrophages was independently validated on the BMDMs harvested from female C57BL/6 mice (**Figure 5F)**. However, the methionine-restricted group showed similar secretory TNF-α levels at both 24 and 48 hpi. In the methionine supplementation group, the probable diversion of carbon flux from serine to one-carbon metabolism was insufficient to drive the pro-inflammatory response, leading to significantly lower IL-1β levels than in the control group at 48 hpi. Methionine supplementation in these BMDMs increased mycobacterial clearance, primarily through its pro-inflammatory effects, as evidenced by elevated extracellular IL-1β and TNF-α levels. Interestingly, the effect of methionine supplementation was abrogated upon treating the macrophages with Caspase-1 inhibitor, leading to lower IL-1β production from Pro IL-1β (**Figure 5G**). This confirmed that the enhanced mycobacterial clearance is a direct attribute of IL-1 response.

### Methionine supplementation has a preventive and therapeutic effect against Mtb in mice

A group of mice received methionine in drinking water (2.5%, 3× times the normal methionine in the diet) along with sucrose (2%) as a vehicle for two weeks. Body weight gain was lower in the supplemented group after the first few doses, becoming similar to that of the control group at later time points (**Supplementary Figures S5A** and **S5B**). The methionine-supplemented group (MS) showed lower water intake but similar food intake compared with the control group (CT) throughout the experiment (**Supplementary Figure S5C**). Random blood glucose levels were similar in both the methionine-supplemented and the control groups (**Supplementary Figure S5D**). Two weeks of methionine supplementation had minimal impact on tissue weights but decreased the proportion and numbers of alveolar macrophages, with an increase in the proportion of non-alveolar monocytic macrophages (**Supplementary Figures S5E, S5F**, **S5G and S5H)**.

We then aerosol-infected the mouse groups with low doses of Mtb infection. We shifted a subgroup of mice from methionine supplementation (MS) to control, which marginally increased the water intake and body weight gain (**Supplementary Figures S6A** and **S6B**). The body weight gain and feed and water intake profiles were consistently similar pre- and post-Mtb infection in the control and supplemented, mouse groups. Methionine supplementation had minimal impact on glucose homeostasis, as we observed similar fasting, random blood glucose, a similar area under the curve after intraperitoneal glucose tolerance test in the Mtb-infected groups (**Supplementary Figure S6C**). The gross tissue pathology, tissue weights (lungs, spleen, and liver), and histopathological scores were similar across all groups (**Supplementary Figures S7A, S7B, and S7C).** However, there was increased glycogen accumulation in the liver and increased germinal centres in the spleen of the methionine-supplemented group (**Supplementary Figure S7D** and **S7E**). S-adenosyl methionine synthase isoform 1 (Mat1), which is primarily expressed in the liver and catalyses the conversion of methionine to SAM, was significantly upregulated in the liver of the methionine-supplemented group compared with infected controls (**Figure 6B**). Both methionine-supplemented groups (continuously supplemented throughout the experiment: MS and supplemented for two weeks pre-infection: MS to CT) showed significantly lower mycobacterial burden in the lungs and spleen at 21 dpi in two independent experiments (**Figure 6C** and **6F**). IL-1β and IFN-γ levels were significantly higher in the lung lysates and serum of the MS groups compared with the CT group (**Figure 6D** and **6E)**. We also observed a higher frequency of Mtb-specific TNF-α+ macrophages in the spleen of the methionine-supplemented groups as compared with CT, and a higher IFN-γ+ activated CD4+ T-cell frequency in the spleen of the methionine-supplemented groups compared with infected controls (**Figure 6F**, **6G**, **6H, Supplementary Figures S8A, S8B** and **S8C**). The MS group showed a significant reduction in activated CD4^+^ T cells in the lungs, with minimal impact on splenic activated CD4+ T cells at 21 dpi (**Supplementary Figure S6D and S6E)**. Since the supplemented groups showed increased IL-1β production, which is mainly secreted by monocytes and macrophages. Increased IL-1β and TNF-α from innate immune cells promote T-cell activation and increased IFN-γ production by T cells. We observed increased IFN-γ in the lungs of methionine-supplemented groups. Interestingly, the Mtb-specific TNF-α-producing macrophages and IFN-γ-producing T-cells, specific against Mtb, showed a higher frequency in the spleens of mice continuously supplemented, as well as when shifted to control post-infection. This showed that 2 weeks of methionine supplementation primed the host’s innate immune cells, resulting in sustained increases in pro-inflammatory cells and better IL-1β levels compared to continuous supplementation, leading to an immunoprophylactic effect.

**Figure 6:**
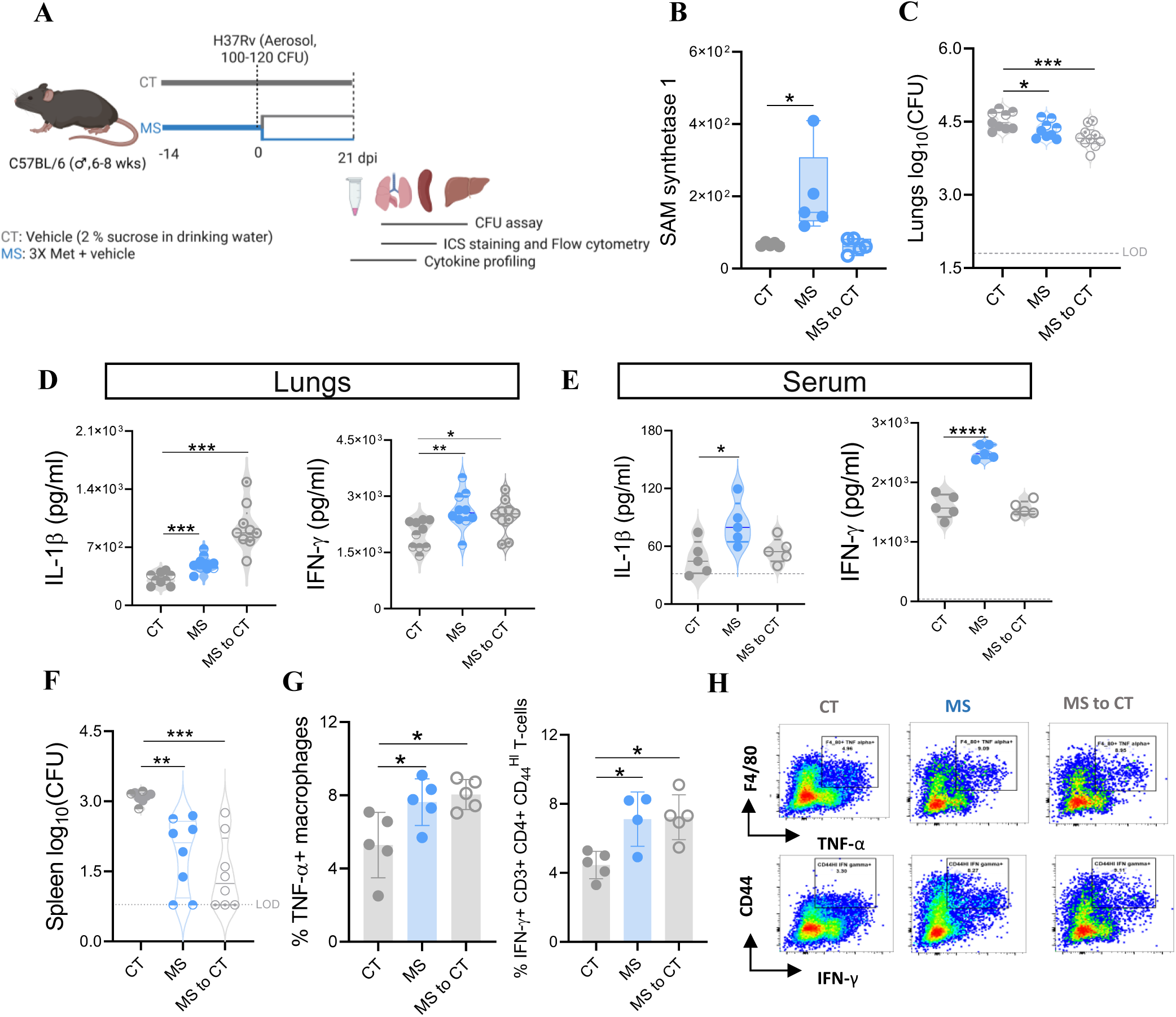
Methionine supplementation improves the inflammatory profile and mycobacterial clearance in the lungs and spleen of Mtb H37Rv-infected mice. **A.** Schematic showing the experimental strategy adopted in this study. **B.** S-adenosyl methionine synthase levels in the liver of the methionine-supplemented mice and controls at 21 dpi. **C.** Mycobacterial burden in the mouse lungs at 21 dpi. **D.** IL-1β and IFN-γ levels in lung lysate. **E.** IL-1β and IFN-γ levels in serum. **F.** Mycobacterial burden in the spleen of methionine-supplemented and control mice at 21 dpi. **G.** Proportions of TNF-α positive macrophages and IFN-γ positive activated CD4+ T-cells upon 6 hours of Mtb-WCL stimulation to the mouse splenocytes harvested at 21 dpi. **H.** Representative plot for the intracellular cytokine staining. One-way ANOVA with Tukey’s multiple comparisons is used, dpi: days post-infection, CT: control, MS: methionine supplemented, MS to CT: Methionine supplemented for 2 weeks, then shifted to control post Mtb infection, WCL: Whole cell lysate, Panel C, D and F include combined data from two independent experiments, dpi: days post-infection. *:p-value<0.05, **:p-value<0.01, ***:p-value<0.001 at 95% confidence.

### Methionine supplementation positively impacts tissue metabolome in Mtb H37Rv-infected mice

Dietary interventions not only impact the gut microbiome but also have a direct impact on tissue metabolome. Hence, we profiled serum, lung, and liver samples from Mtb-infected mice supplemented with methionine and controls, and amino acid levels showed significant variation (**Supplementary Figure S9A**). Absolute quantification using a standard curve showed that methionine levels were significantly higher in serum and lungs; however, the absolute amounts were within the normal range. Homocysteine levels were similar and within the normal range across all three tissues, indicating that the methionine intervention was not toxic (**Figure 7**). In circulation, we observed 723 putatively annotated metabolic features, out of which 20 (10/10: higher/lower) showed significant alterations in the methionine-supplemented group. Pathway enrichment analysis of the deregulated features showed methionine, serine, polyamine and nucleotide metabolism to be enriched based on upregulated features, and tryptophan metabolism showed downregulation (**Supplementary Figure S9B**). Tryptophan metabolites, especially kynurenine, produced by the indoleamine 2,3-dioxygenase (IDO) enzyme, are reported to suppress host immune response against pathogens. (Gautam et al., 2018) Hence, decreased kynurenine levels in the methionine-supplemented group mice’s serum would result in lesser suppression and hence better immune response. The lung and liver metabolomes showed downregulation of lipid biosynthesis and enrichment of polyamine and methionine metabolism (**Figure 7** and **Supplementary Figure S9B**). Enhanced lipid biogenesis is well reported to promote bacterial growth in the liver and lung tissue. (Sankar et al., 2024)(Sarkar et al., 2025) Methionine is reported to increase the expression of lipid efflux transporters, which leads to reduced lipid accumulation and synthesis in the methionine-supplemented group. The liver of the methionine-supplemented mice group showed increased arginine biosynthesis, which is crucial for reducing toxicity and improving liver profile upon drug interventions. (Dong et al., 2024) However, we did not observe any correlation between and spleen metabolite levels, suggesting a targeted impairment in lung tissue (**Supplementary Figure S9C**). Hence, methionine supplementation not only supports amino acid metabolism and pro-inflammatory response but also positively impacts host lung, liver and circulatory metabolite profiles.

**Figure 7:**
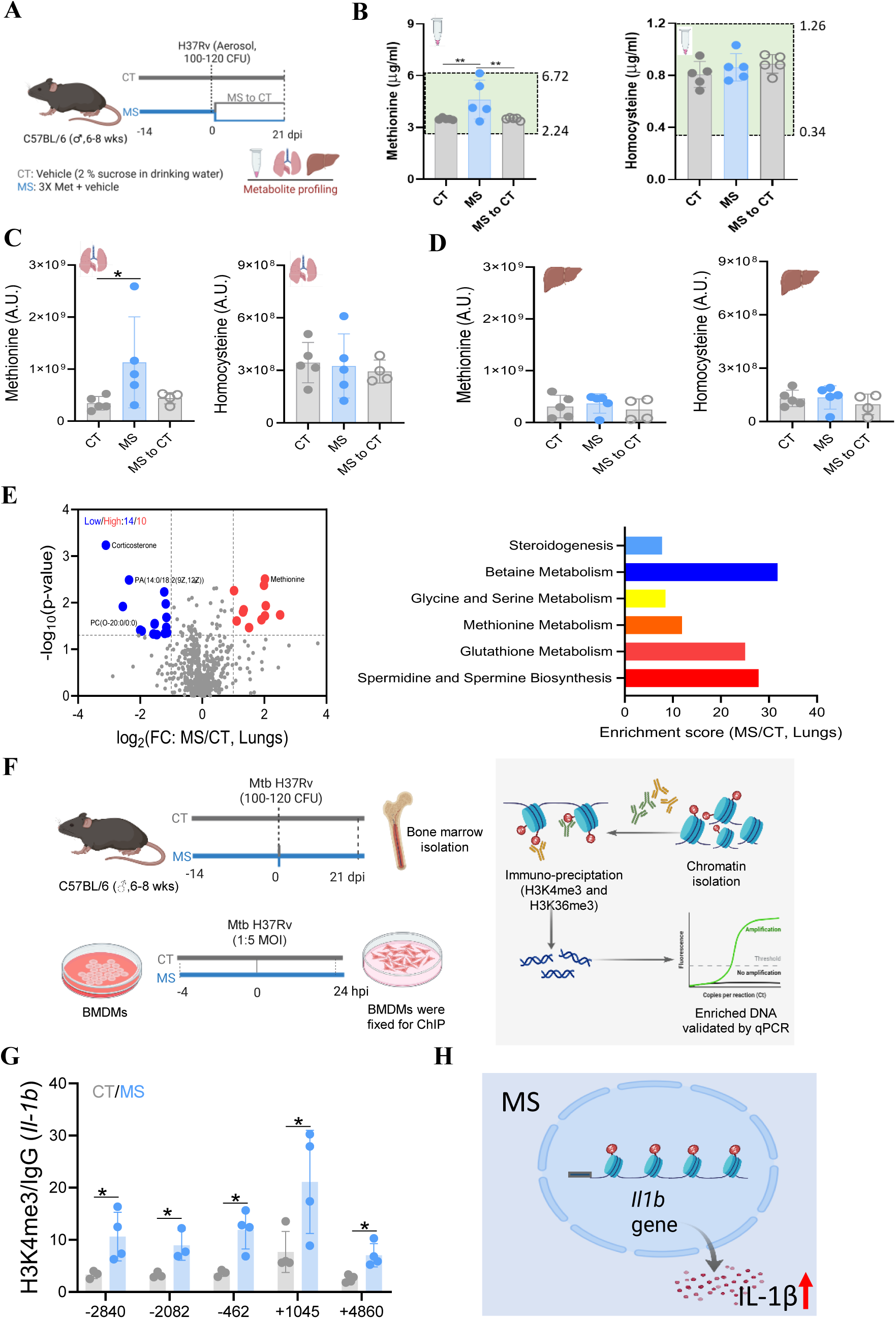
Methionine supplementation improves lung metabolic profile in Mtb H37Rv-infected mice and increases H3K4me3 chromatin occupancy for *Il-1b* gene. **A**. Schematic showing the experimental strategy adopted in this study. **B.** Absolute quantification of methionine and homocysteine in the serum of various mouse groups, where the green box shows the normal amount of methionine and homocysteine in healthy mouse serum. **C** and **D.** Relative quantification of methionine and homocysteine in lungs and liver, respectively. **E.** Volcano plot showing the significantly altered metabolic features and the pathway enrichment plots for lungs of methionine-supplemented mice compared with infected controls. **F.** Schematic showing the experimental strategy adopted in the study. **G** and **H.** ChIP qPCR of H3K4me3 and H3K36me3 enrichment at different positions for *Il-1b* gene, respectively. **D.** Methionine supplementation increases IL-1β production by modulating histone methylation in Mtb-infected macrophages. One-way ANOVA with Tukey’s multiple comparisons is used; log_2_FC > ±1.0 and log10 (p-value)> 1.3 are considered significant for differences; dpi: days post-infection. *:p-value< 0.05, **:p-value<0.01 at 95% confidence. The green box represents the normal range for methionine and homocysteine levels in healthy mouse serum.

### Methionine supplementation affects the liver proteome with improved NF-kB and decreased PPAR signalling in Mtb-H37Rv-infected mice

We performed proteomics analysis of the liver of methionine-supplemented and control mice groups, which showed distinct clusters in the partial least squares-discriminant analysis (PLS-DA) plot (**Supplementary Figure S10A**). A set of 1996 proteins was identified; 79 proteins (35/44: down/up) were deregulated in the methionine-supplemented group compared with the infected control at 21 dpi (**Supplementary Figure S10B**). Pathway enrichment analysis showed increased translation of proteins, enhanced detoxification, and improved response to oxidative stress, along with downregulation of cholesterol biosynthesis. Shifting mice from methionine supplementation to control upon infection also showed a distinct profile with 65 deregulated proteins (18/47: down/up) compared with infected controls. The MS to CT group showed increased translation, improved pro-inflammatory response via TNF-α/NF-kB signalling and downregulated PPAR signalling (**Supplementary Figure S10C**). However, this group showed similar levels of Mat1, emphasizing that SAM production is not the only way methionine supplementation improves bacterial clearance. We observed significant upregulation of kynurenine hydroxylase, which shifts metabolites from tryptophan metabolism towards ROS generation and an anti-microbial environment, and glutathione reductase, which is important for preventing excessive inflammation-associated tissue damage by detoxifying ROS. Overall, liver proteomics data indicate improved immune response and increased detoxification in the mice receiving methionine supplementation.

### Methionine supplementation improves the antioxidant potential of the Mtb-H37Rv-infected host

We performed elemental analysis of the serum from methionine-supplemented and control mice using ICP-MS (**Supplementary Figure S11**). Methionine-supplemented and crossover group mice showed a significant increase in Selenium (Se) levels compared with infected controls at 21 dpi (**Supplementary Figure S11A**). Iron (Fe) and magnesium (Mg) showed marginal differences, whereas calcium (Ca) showed a significant increase in the methionine-supplemented group, which abrogated upon switching the mice to control water (**Supplementary Figures S11B, S11C** and **S11D**). The copper (Cu) to zinc (Zn) ratio, which determines the antioxidant potential of the host, showed a significant decrease in the methionine-supplemented mice, whereas Cu showed a significant decrease in the crossover mice, together indicating improved antioxidant capacity of the host (**Supplementary Figures S11E, S11F** and **S11G**). We performed ELISA to monitor SOD and observed a significant increase in SOD activity in the methionine-supplemented and crossover groups compared with infected controls at 21 dpi (**Supplementary Figure S11H**). Interestingly, the MS-to-CT group showed significantly higher SOD activity than the MS group, suggesting much greater antioxidant potential. Hence, methionine supplementation also improves the overall antioxidant potential of the Mtb-infected host.

### Methionine supplementation trains bone marrow precursors for improved immune response

From the *in vivo* methionine-supplemented and control mice, we harvested bone marrow and differentiated them into macrophages in either methionine-supplemented or control media (**Supplementary Figure S12A**). We observed reduced bacterial burden in Mtb-infected BMDMs from *ex vivo-*supplemented (CT-exMS) macrophages, as well as *in vivo*-supplemented macrophages differentiated in supplemented media (MS-exMS; **Supplementary Figure S12B**). Interestingly, the effect of methionine supplementation persisted even after discontinuing methionine supplementation during macrophage differentiation (MS-exCT), as evidenced by a lower bacterial burden compared with the control group. IL-1β and TNF-α production were significantly higher in the MS-exMS group as compared with the control (**Supplementary Figure S12C**). Methionine supplementation increased the pro-inflammatory response in the lungs, thereby enhancing bacterial clearance in the Mtb-infected mice. Methionine supplementation in mice reprograms bone marrow precursors, thereby improving mycobacterial clearance in an *ex vivo* Mtb infection model.

### Methionine supplementation increases expression of the *Il-1b* gene by enhancing H3K4 trimethylation in BMDMs

Methionine supplementation increases the pro-inflammatory response upon Mtb infection in mice by enhancing IL-1β and TNF-α production. Several histone H3 methylation marks, including trimethylation at lysines 4 and 36 (H3K4me3 and H3K36me3), can have a substantial role in defining chromatin state and regulating inflammatory gene expression. (Shailatifard, 2006) We used ChIP coupled with qPCR to investigate how H3K36me3 and H3K4me3 chromatin occupancy of specific inflammatory marker genes (*Il-1b* and *Tnf-a*) changed upon methionine supplementation in Mtb-infected BMDMs at 24 hpi. We observed increased enrichment of H3K4me3 and H3K36me3 at the *Il-1b* promoter by ChIP-qPCR. However, H3K4me3 enrichment at *Il-1b* was significantly higher in Mtb-infected methionine-supplemented macrophages at 24 hpi as compared to infected controls (**Figure 7F**, **7G** and **7H**). Since we did not observe increased H3K4me3 or H3K36me3 enrichment at the *Tnf-a* gene, the increased TNF-α levels in our data seem to be a direct consequence of an increase in the pro-inflammatory profile of macrophages upon methionine supplementation rather than direct methylation of the gene promoter (**Supplementary Figure S12D**). These findings provide direct evidence for a dynamic regulatory role of methionine via histone methylation at critical amino acid residues that mediate cellular epigenetic status and regulate gene expression to enhance the pro-inflammatory response against Mtb.

## Discussion

Cellular metabolism is an important contributor to regulating immune cell function. (Caputa et al., 2019)(Muri & Kopf, 2021) While the spotlight in recent years has been on understanding central carbon metabolism, few studies have examined the contribution of one-carbon metabolism to the compromised functionality of Mtb-infected macrophages. (Wculek et al., 2022)(Pagán et al., 2022)(Woods et al., 2020) One carbon metabolism generates S-adenosyl methionine (SAM), which transfers carbon units for histone methylation and regulates the transcription of genes related to pro-inflammatory cytokines, such as IL-1β, upon LPS or IFN-γ stimulation. (Yu et al., 2019)(Wang et al., 2024) Methionine and serine are the primary contributors to the carbon units for one-carbon metabolism. Immune cells like macrophages can synthesise serine via the glycine cleavage system, whereas methionine, being an essential amino acid, needs to be sourced extracellularly. We observed decreased expression of *Mthfr, Mthfd,* and higher *Slc7a5, Msra, Msrb, Phgdh* expression in the alveolar macrophages, highlighting decreased one-carbon pool and methionine levels in the cells, leading to higher salvage. Alveolar macrophages were unable to undergo the infection-induced metabolic switch to glycolysis needed for a pro-inflammatory phenotype. These findings suggest that downregulation of one-carbon metabolism and impaired metabolic switching may altered IL-1 responses, thereby promoting bacterial survival in alveolar macrophages.

Next, we aimed to understand the early metabolic changes in the *ex vivo* Mtb-infected macrophages. The metabolic perturbations in Mtb-infected BMDMs showed a major shift at 24 hpi, with downregulation of various nodes of one-carbon metabolism. Serine-methionine metabolism and polyamine synthesis were downregulated in these BMDMs. However, the nucleotide salvage pathway was significantly upregulated upon Mtb infection. Significantly higher expression of IL-1 and caspase was also observed at 0, and 4 hpi, leading to controlled bacterial burden in the early phases. However, bacterial clearance was dampened at 24 hpi with a significantly higher mycobacterial burden and decreased intracellular serine and methionine levels. We observed significant downregulation of IL-1 and caspase-1 levels from 0 to 24 hpi in BMDMs. Our findings suggest that decreased methionine and serine, and increased nucleotides like adenine, contribute to the suppressed immune response at 24 hpi in Mtb-infected macrophages. Interestingly, glutathione, produced from methionine via the transsulfuration pathway, showed similar levels at early time points and significantly increased at 24 hpi in Mtb-infected macrophages. The delay in increasing glutathione production might be helping the bacteria establish a favourable niche and replicate early, leading to an increased bacterial burden at 24 hpi. It would be interesting to see whether the decreased levels of serine, cysteine, glutamine, and glycine are partly contributing to the observed higher tripeptide glutathione levels. Even though arginase expression was lower in Mtb-infected macrophages, diverting the flux towards nitric oxide synthesis was insufficient to control bacterial growth.

Since methionine acts as a functional link between epigenetic and metabolic reprogramming of immune response (Daskalaki et al., 2025), we confirmed the diversion of carbon units from methionine of Mtb-infected BMDMs. In our stable isotope labelling experiment using ^13^C-Methionine in culture, we observed the diversion of carbon units towards the polyamine metabolism node, which supports nucleotide salvage. Hence, we report that the extracellular methionine pool is diverted toward polyamine synthesis, leading to increased nucleotide salvage and a reduced carbon pool for methylation. Hence, we hypothesised that supplementing appropriate metabolites, such as methionine, would help rescue macrophage function by increasing the available carbon units for methylation.

Dietary methionine restriction is reported to suppress inflammation in various immune cells, such as macrophages and T cells, whereas supplementation is reported to improve macrophage function. (Roy et al., 2020)(Franceschi et al., 2020)(Santos et al., 2017)(W.-W. Zhang, Xiang, et al., 2024) BMDMs supplemented with 1 mM methionine (i.e. 5 times the normal concentration) upon Mtb infection showed improved cell viability and a lower mycobacterial burden. Methionine supplementation significantly increased IL-1β production in Mtb-infected macrophages, which might contribute to higher phagocytosis, as evidenced by lower bacterial burden in the supplemented group. Inflammasome activation by IL-1β and phagolysosome fusion by TNF-α might contribute to the increased bacterial clearance. (Olsen et al., 2016)

A sustained pro-inflammatory response damages host tissues, and supplementing macrophages with methionine increased their acute pro-inflammatory cytokine production compared to control at both 24 and 48 hpi. However, there was a drop at 48 hpi, allowing the host cells to control the response and limiting tissue damage. Methionine is a positive regulator of cholesterol efflux by increasing ABCA1 and LXR-α expression. (O’Callaghan et al., 2022) Mtb infection increases lipid accumulation, leading to the formation of foamy macrophages. (Hadchouel et al., 2022) Even though methionine supplementation is reported not only to enhance pro-inflammatory responses in immune cells but also to improve cognitive function, gut health, lifespan, reduce the risk of colon cancer, and promote haematopoiesis. (Xu et al., 2024)(W.-W. Zhang et al., 2024)(Miousse et al., 2017)(Fanti et al., 2026) However, supplementation at higher amounts (>6×) of methionine or for a longer duration can have an adverse impact on the host as it leads to homocysteine and methanethiol-cysteine disulfides production. (Ishii et al., 2022)

We supplemented drinking water with methionine (2.5%, i.e., 3 times the normal methionine level in the diet) and sucrose to increase palatability. The concentration up to 6 times the normal methionine amount in the mouse diet is reported to be safe, to improve cognitive function, haematopoiesis, and T-cell activation, and to not cause any adverse effects, including hyperhomocysteinemia. (Ishii et al., 2022)(Ligthart-Melis et al., 2020)(De Rezende & D’Almeida, 2014)(Xu et al., 2024)(Miousse et al., 2017) Methionine amount in diet also regulates glucose homeostasis, and restricting it is reported to improve glucose homeostasis. (Ishii et al., 2022)(Y. Zhang et al., 2024) Methionine along with vitamin B complex is also reported to reduces anti-Tb drugs associated liver toxicity.(Amagon et al., 2017) Methionine supplementation had minimal impact on body weight gain, feed intake, glucose homeostasis and tissue weight but impacted water intake due to the sulphurous smell. At 3× dosages, methionine levels were significantly increased in circulation as well as the liver without any toxic homocysteine buildup. Two weeks of methionine supplementation improved the lung inflammatory profile by decreasing the proportion of anti-inflammatory alveolar macrophages and increasing the proportion of the pro-inflammatory monocytic macrophages. After 2 weeks of supplementation, these mice were infected with Mtb, and a subset was shifted to a vehicle control. We hypothesised that 2 weeks of methionine supplementation would be sufficient to reprogram immune cells and improve immune response. Methionine supplementation is reported to improve alveolar macrophage function by promoting a pro-inflammatory response, lipid efflux, and T-cell function, while preventing exhaustion. (Sharma et al., 2025) (O’Callaghan et al., 2022) We also observed decreased lipid biogenesis in the lungs and liver of the methionine-supplemented group, accompanied by beneficial metabolic changes, such as increased SAM production, increased arginine biosynthesis in the liver, and decreased tryptophan metabolism in circulation. Methionine-supplemented groups showed enhanced mycobacterial clearance in the lungs and spleen at 21 dpi. Methionine supplementation also trained bone marrow cells to mount a stronger immune response and improved mycobacterial clearance. This effect persisted even after discontinuing supplementation, and upon differentiation of the BMDMs, a reduced mycobacterial burden was observed. Methionine supplementation does not unilaterally increase inflammation, but it has a balanced effect on the regulatory anti-oxidative response and mitigates inflammation-associated tissue damage in the host. (Troha et al., 2026) We also observed increased levels of Mg and Ca in the systemic circulation. Mg protects mitochondria from oxidative stress, and Ca helps shift macrophages toward M1 polarisation and increases their pro-inflammatory function, thereby providing a balanced inflammatory response. (Sun et al., 2020)(Seegren et al., 2023)(Pogonyalova et al., 2025) It is well reported that methionine supplementation increases pro-inflammatory response at the site of infection; however, it also increases the release of excess cytokines via the kidneys and improves renal function. (Troha et al., 2026) However, our study did not focus on renal clearance; we did observe a significant increase in the host’s overall antioxidant capacity. Total Superoxide dismutase activity was significantly higher in the serum of methionine-supplemented mice groups compared with vehicle controls. Also, selenium levels were higher, and the copper-to-zinc ratio was lower. Selenium serves as the cofactor for glutathione peroxidase (GPx) and enhances the antioxidant capacity of cells. (Nelson et al., 2011)(Vunta et al., 2007) Cu is needed for Superoxide dismutase activity and indicates elevated oxidative stress; Zn inhibits ROS production. (Stafford et al., 2013) A lower Cu or a lower Cu-to-Zn ratio indicates better antioxidant potential in the Mtb-infected methionine-supplemented mice.

Methionine supplementation directly affects SAM production. SAM act as a methylation substrate for DNA and histone methylation reactions. Ultimately, changes in the histone modification state can contribute to altered cytokine expression. Several histone H3 methylation marks, including those at lysines 4 and 36 (H3K4me3 and H3K36me3), play a substantial role in defining chromatin state and regulating gene expression. (Shilatifard, 2006) Genome-wide assessment of the H3K4me3 mark suggested that cells under methionine restriction attempted to maintain SAM levels by decreasing its consumption and using alternative mechanisms to maintain the H3K4me3 mark. Alterations in H3K4me3 directly feeds back to one-carbon metabolism and methylation-related enzymes to maintain physiological homeostasis. (Mentch et al., 2015) It is well established that high SAM: SAH ratios play an important role in maintaining histone methylation reactions, such as H3K4me3 and H3K36me3, during LPS-induced inflammation in peritoneal macrophages.(Yu et al., 2019) As in the above reports, we also observed enrichment of H3K4me3 and H3K36me3 at the promoters of *Il-1b* and *Tnf-a* by ChIP-qPCR. In our study, H3K4me3 enrichment at *Il-1b* was significantly increased at 24 hpi in the methionine-supplemented group compared with Mtb-infected control macrophages. Therefore, as an essential amino acid, exogenous methionine serves as the primary methyl donor for SAM generation and subsequent methylation reactions, which are essential for Mtb-induced IL-1β production.

In conclusion, dietary methionine supplementation improves macrophage function, leading to increased production and secretion of IL-1β, which can contribute to an acute pro-inflammatory response and increased T-cell activation, thereby improving clinical outcomes during chronic Mtb infection.

### Limitations of the study

The priming of T cells by methionine-supplemented macrophages to enhance IFN-γ production needs further investigation. We observed both immunoprophylactic and therapeutic benefits of methionine supplementation in Mtb-infected mice; however, it will be interesting to determine whether this supplementation strategy could serve as an adjunct to existing anti-TB drugs to improve clinical outcomes.

## Online supplemental material

Supplemental Figures are available in the Supplemental File.

## Acknowledgements

We acknowledge the Department of Biotechnology (DBT), Government of India, for supporting activities through research grants and supporting the Tuberculosis Aerosol Challenge Facility at ICGEB and ICGEB New Delhi, which provided core support to RKN. Nidhi Yadav, Ashish Gupta and Mothe Sravya received a Junior Research Fellowship from the DBT. Satya Ranjan Sahoo and Suchitra Jena received a Junior Research Fellowship from Council of Scientific and Industrial Research - University Grants Commission. Nainy Goel is supported by Ministry of Earth Sciences (MoES) project. We acknowledge Dr Anmol Chandele for her valuable contributions throughout the study. We acknowledge Suraj Nayar for the help with Compound Discoverer-based mass spectrometry data analysis. We acknowledge Shweta Chaudhary for sharing the raw scRNAseq experimental data for reanalysis. We acknowledge Dr Tenzin and Suneet from the LC-MS facility at ICGEB, New Delhi, for their help with proteomics data acquisition. We acknowledge Dr Vivek Rao from IGIB for kindly sharing reagents. We acknowledge the BCIL-IP cell for assistance in filing a provisional patent under Indian patent application no. 202511081045.

## Authorship Contributions

NY and RKN conceptualised and designed experiments; NY, AG, SRS, SJ, and NG performed the experiments; NY analysed the data and prepared figures. NS and JSM helped with the LC-MS/MS data acquisition. MS, TN and DD helped with scRNAseq data analysis. SNJ and NG helped with flow cytometry data acquisition. BNP carried out the histopathological scoring. SKM and AKP shared expertise in the data analysis and reagents. RKN supervised the study and did formal analysis. Funds for this work were generated by RKN; NY and RKN wrote the first draft of the manuscript and revised it, incorporating the comments of all co-authors.

## Abbreviations

TB: Tuberculosis
BM: Bone marrow
BMDM: Bone marrow-derived macrophages
Mtb: Mycobacterium tuberculosis
CFU: Colony forming units
DMEM: Dulbecco’s modified eagle medium
FBS: Fetal bovine serum
MOI: Multiplicity of infection

