## Supplementary File for "Methionine modulates the metabolic-epigenetic axis as immunotherapeutic in tuberculosis"

### Slide 1
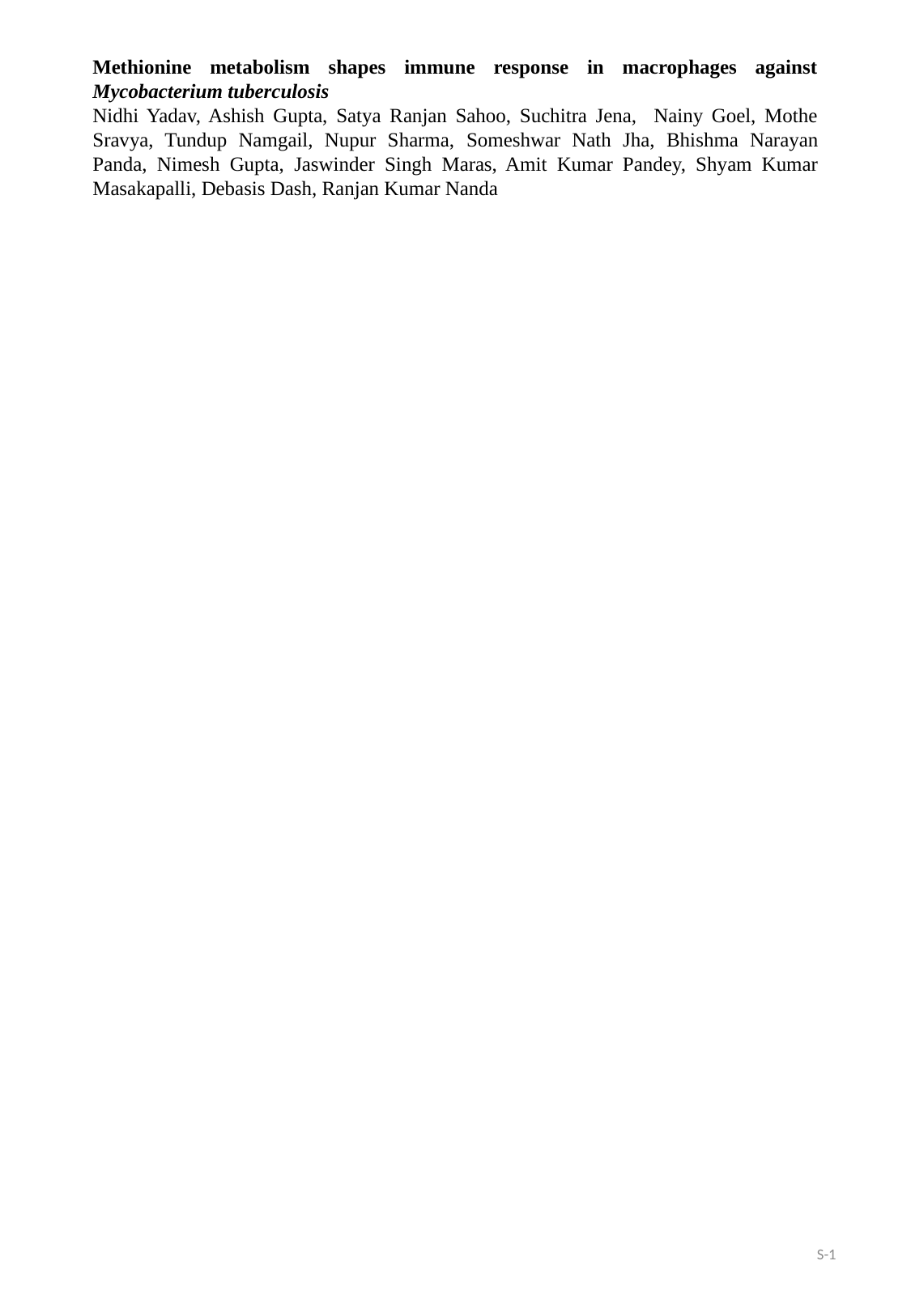

Methionine metabolism shapes immune response in macrophages against Mycobacterium tuberculosis
Nidhi Yadav, Ashish Gupta, Satya Ranjan Sahoo, Suchitra Jena, Nainy Goel, Mothe Sravya, Tundup Namgail, Nupur Sharma, Someshwar Nath Jha, Bhishma Narayan Panda, Nimesh Gupta, Jaswinder Singh Maras, Amit Kumar Pandey, Shyam Kumar Masakapalli, Debasis Dash, Ranjan Kumar Nanda
S-1

### Slide 2
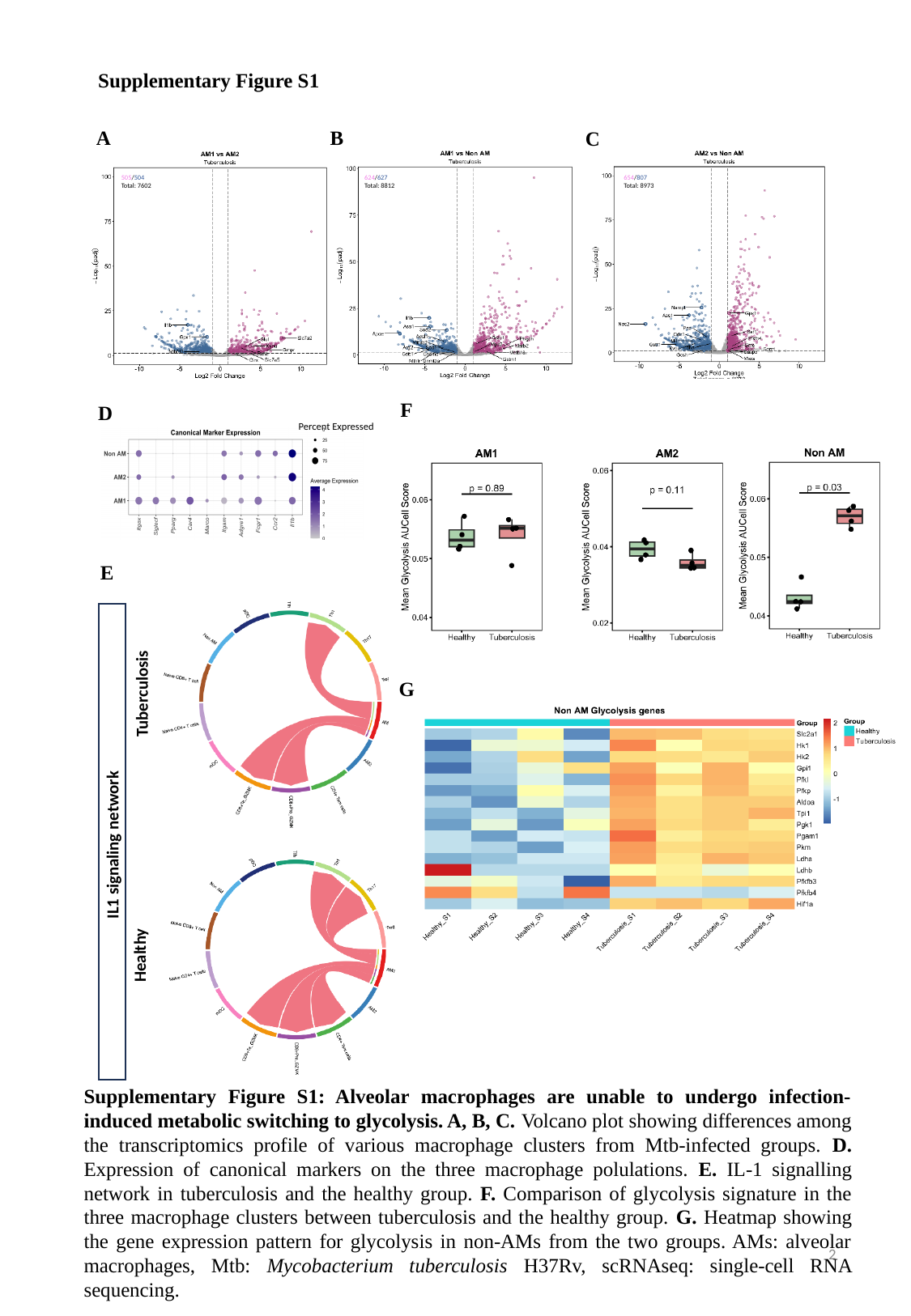

Supplementary Figure S1
A
B
C
654/807
Total: 8973
624/627
Total: 8812
505/504
Total: 7602
F
D
Percent Expressed
E
IL1 signaling network
Healthy
Tuberculosis
G
Supplementary Figure S1: Alveolar macrophages are unable to undergo infection-induced metabolic switching to glycolysis. A, B, C. Volcano plot showing differences among the transcriptomics profile of various macrophage clusters from Mtb-infected groups. D. Expression of canonical markers on the three macrophage polulations. E. IL-1 signalling network in tuberculosis and the healthy group. F. Comparison of glycolysis signature in the three macrophage clusters between tuberculosis and the healthy group. G. Heatmap showing the gene expression pattern for glycolysis in non-AMs from the two groups. AMs: alveolar macrophages, Mtb: Mycobacterium tuberculosis H37Rv, scRNAseq: single-cell RNA sequencing.
2

### Slide 3
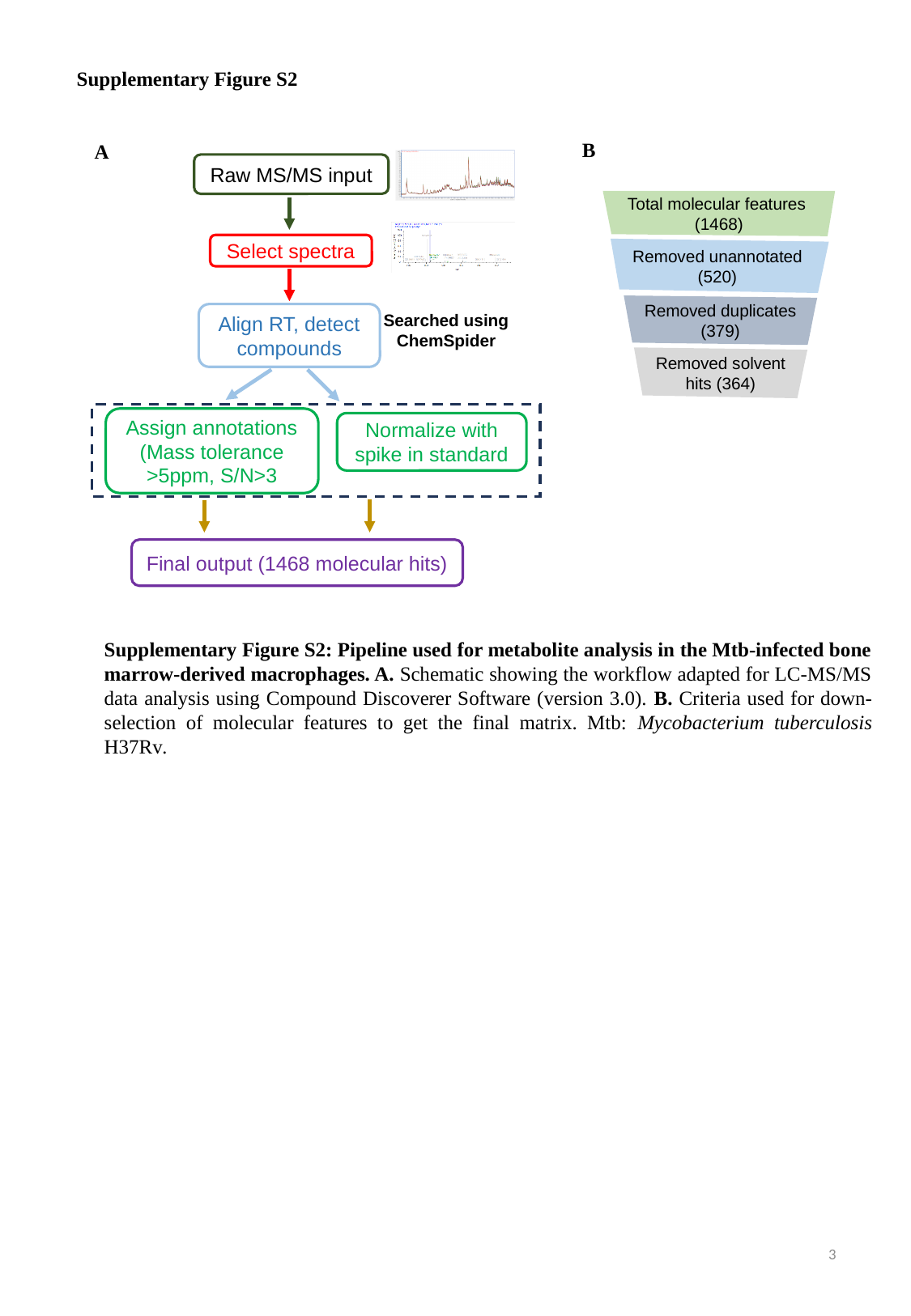

Supplementary Figure S2
B
A
Raw MS/MS input
Select spectra
Align RT, detect compounds
Searched using ChemSpider
Assign annotations
(Mass tolerance
>5ppm, S/N>3
Normalize with spike in standard
Final output (1468 molecular hits)
Total molecular features (1468)
Removed unannotated
(520)
Removed duplicates (379)
Removed solvent hits (364)
Supplementary Figure S2: Pipeline used for metabolite analysis in the Mtb-infected bone marrow-derived macrophages. A. Schematic showing the workflow adapted for LC-MS/MS data analysis using Compound Discoverer Software (version 3.0). B. Criteria used for down-selection of molecular features to get the final matrix. Mtb: Mycobacterium tuberculosis H37Rv.
3

### Slide 4
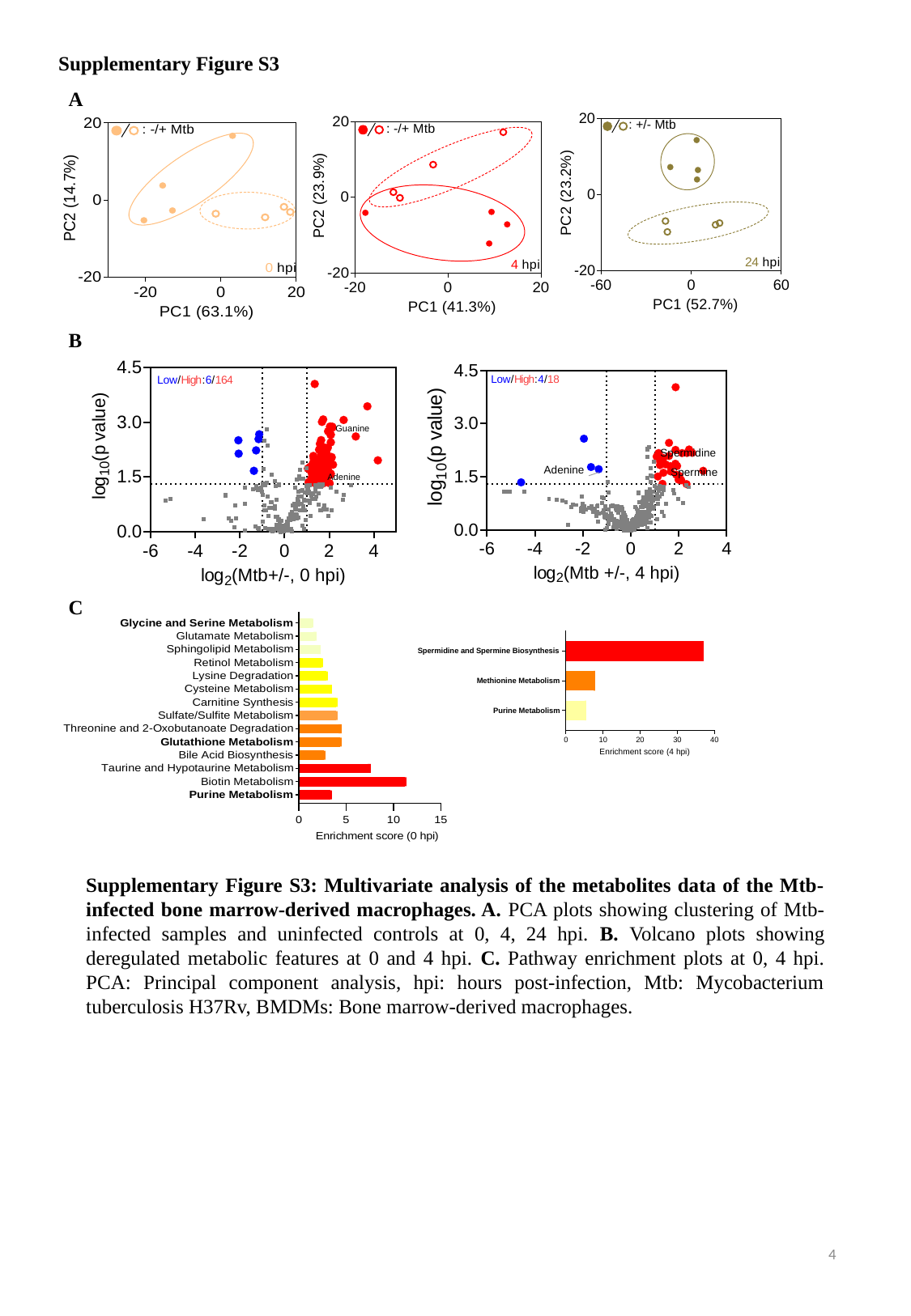

Supplementary Figure S3
A
B
C
Supplementary Figure S3: Multivariate analysis of the metabolites data of the Mtb-infected bone marrow-derived macrophages. A. PCA plots showing clustering of Mtb-infected samples and uninfected controls at 0, 4, 24 hpi. B. Volcano plots showing deregulated metabolic features at 0 and 4 hpi. C. Pathway enrichment plots at 0, 4 hpi. PCA: Principal component analysis, hpi: hours post-infection, Mtb: Mycobacterium tuberculosis H37Rv, BMDMs: Bone marrow-derived macrophages.
4

### Slide 5
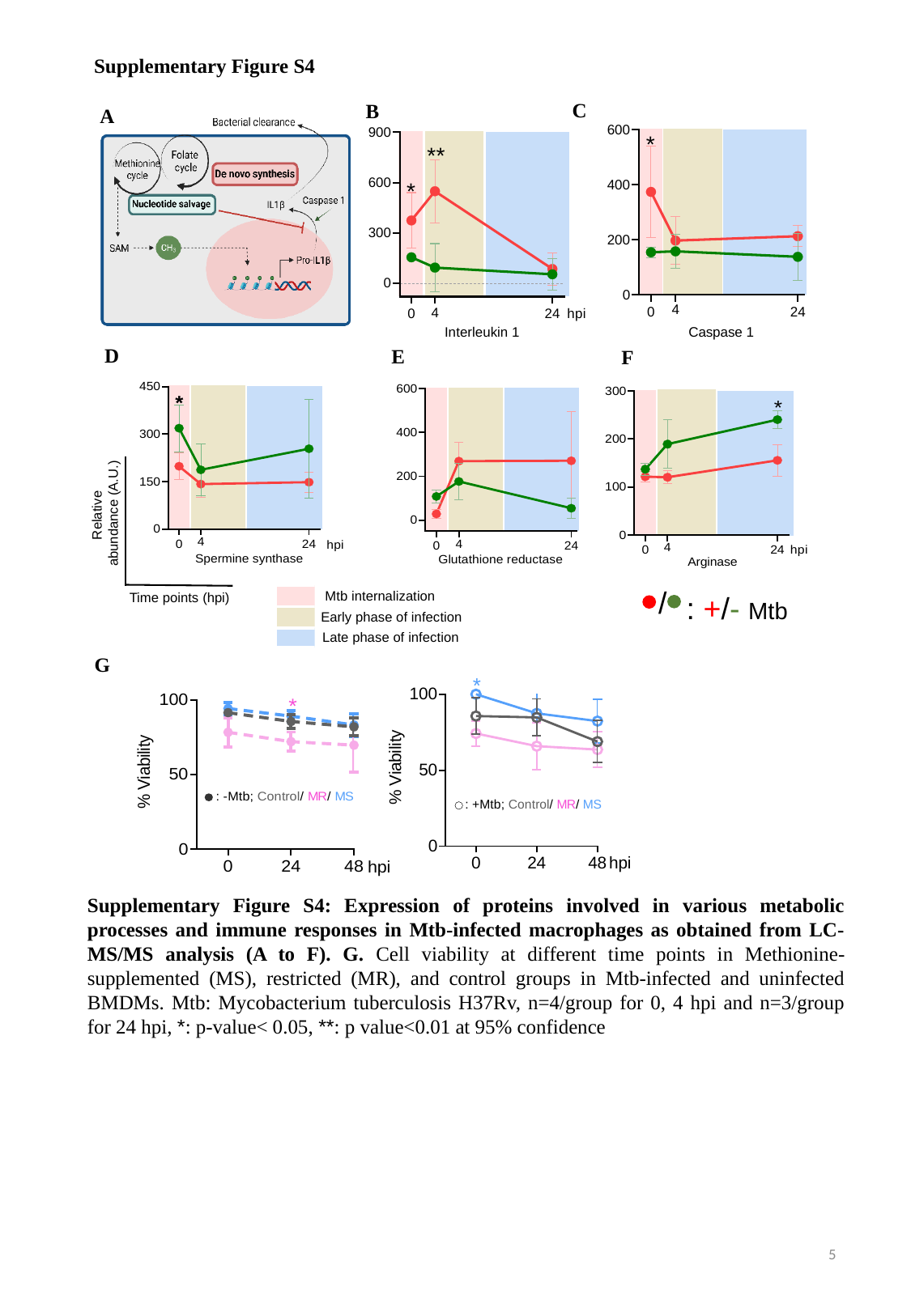

Supplementary Figure S4
C
B
A
D
E
F
Relative
 abundance (A.U.)
Time points (hpi)
: +/- Mtb
/
Mtb internalization
Early phase of infection
Late phase of infection
G
Supplementary Figure S4: Expression of proteins involved in various metabolic processes and immune responses in Mtb-infected macrophages as obtained from LC-MS/MS analysis (A to F). G. Cell viability at different time points in Methionine-supplemented (MS), restricted (MR), and control groups in Mtb-infected and uninfected BMDMs. Mtb: Mycobacterium tuberculosis H37Rv, n=4/group for 0, 4 hpi and n=3/group for 24 hpi, *: p-value< 0.05, **: p value<0.01 at 95% confidence
5

### Slide 6
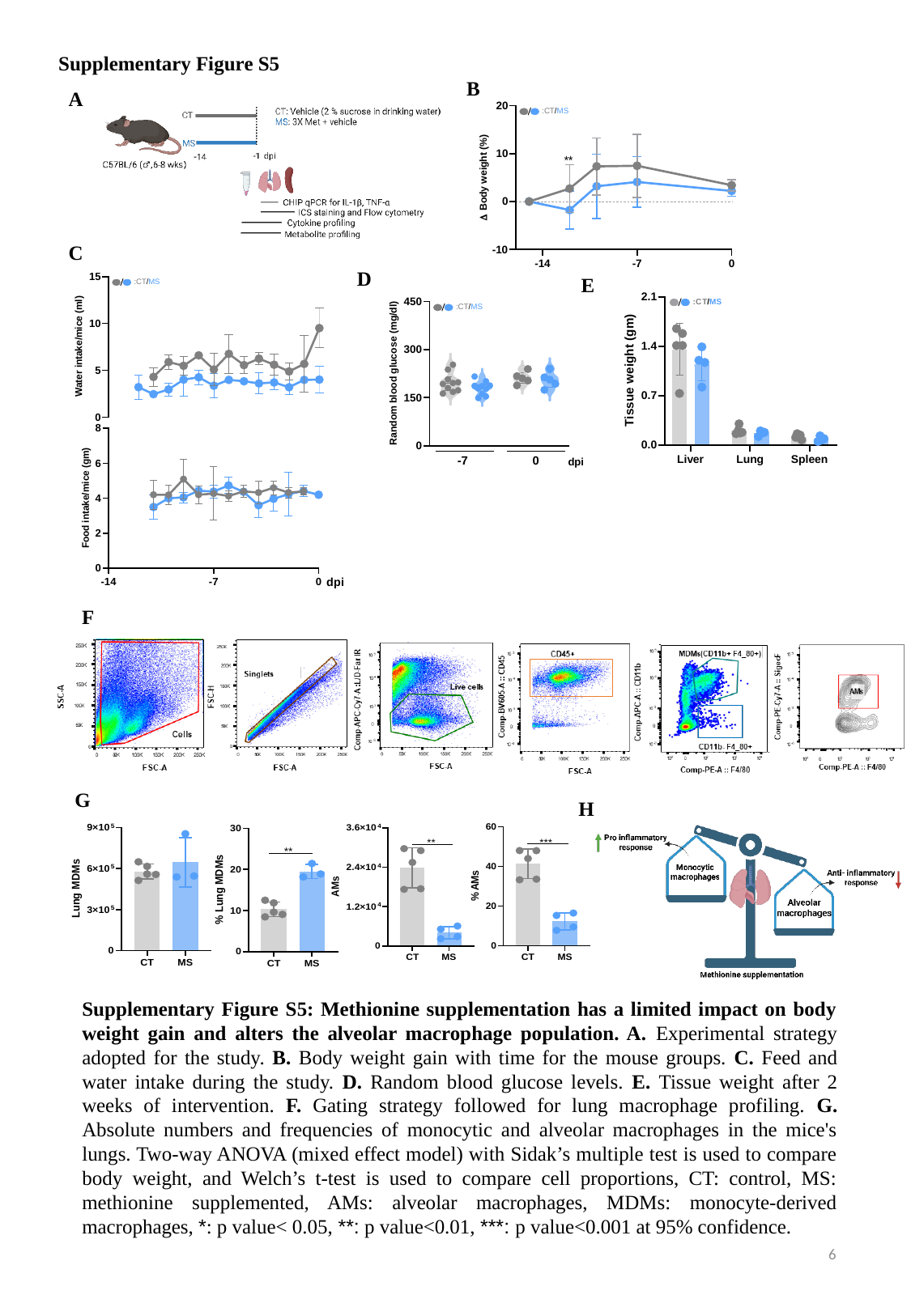

Supplementary Figure S5
B
A
C
D
E
F
G
H
Supplementary Figure S5: Methionine supplementation has a limited impact on body weight gain and alters the alveolar macrophage population. A. Experimental strategy adopted for the study. B. Body weight gain with time for the mouse groups. C. Feed and water intake during the study. D. Random blood glucose levels. E. Tissue weight after 2 weeks of intervention. F. Gating strategy followed for lung macrophage profiling. G. Absolute numbers and frequencies of monocytic and alveolar macrophages in the mice's lungs. Two-way ANOVA (mixed effect model) with Sidak’s multiple test is used to compare body weight, and Welch’s t-test is used to compare cell proportions, CT: control, MS: methionine supplemented, AMs: alveolar macrophages, MDMs: monocyte-derived macrophages, *: p value< 0.05, **: p value<0.01, ***: p value<0.001 at 95% confidence.
6

### Slide 7
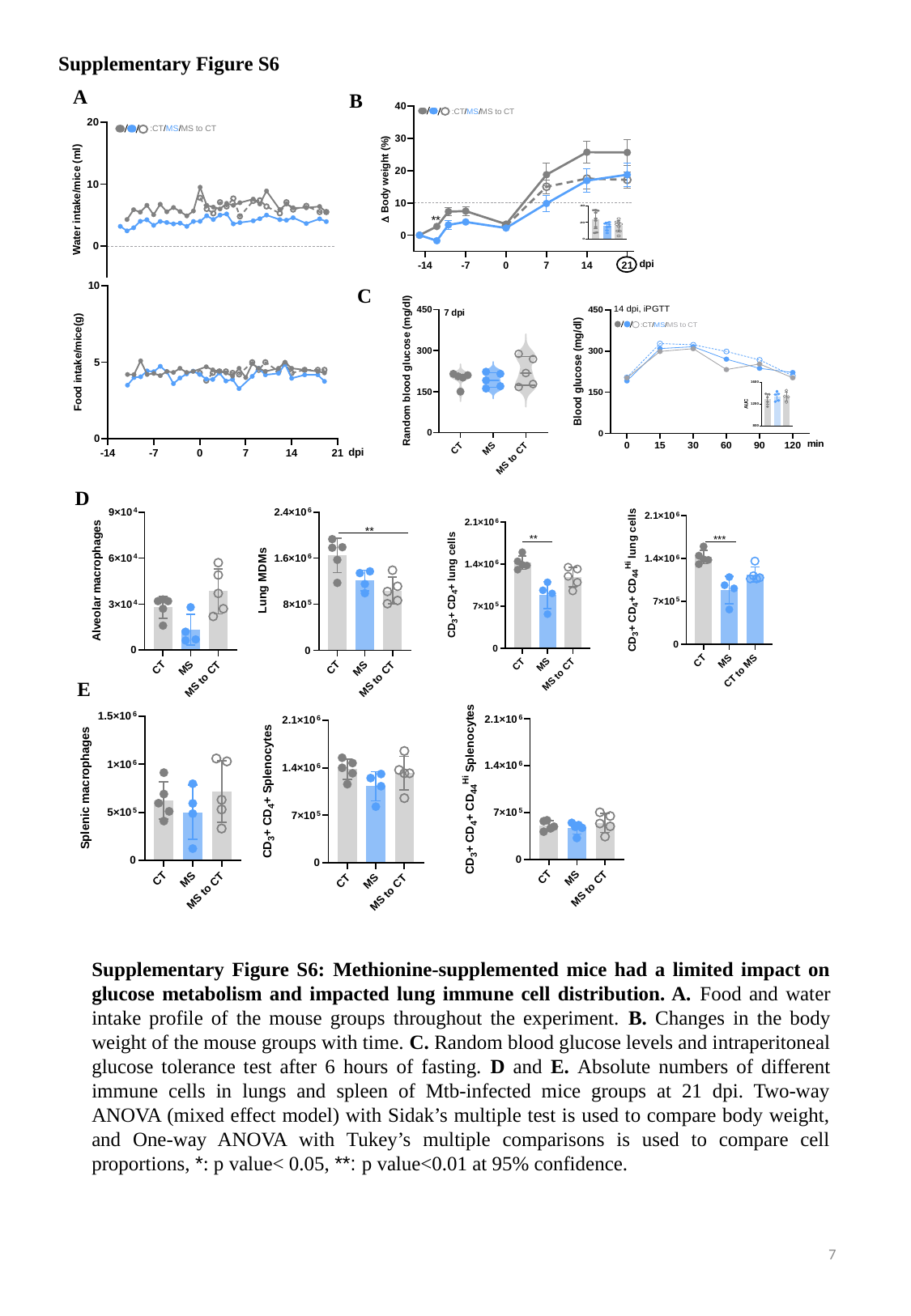

Supplementary Figure S6
A
B
C
D
E
Supplementary Figure S6: Methionine-supplemented mice had a limited impact on glucose metabolism and impacted lung immune cell distribution. A. Food and water intake profile of the mouse groups throughout the experiment. B. Changes in the body weight of the mouse groups with time. C. Random blood glucose levels and intraperitoneal glucose tolerance test after 6 hours of fasting. D and E. Absolute numbers of different immune cells in lungs and spleen of Mtb-infected mice groups at 21 dpi. Two-way ANOVA (mixed effect model) with Sidak’s multiple test is used to compare body weight, and One-way ANOVA with Tukey’s multiple comparisons is used to compare cell proportions, *: p value< 0.05, **: p value<0.01 at 95% confidence.
7

### Slide 8
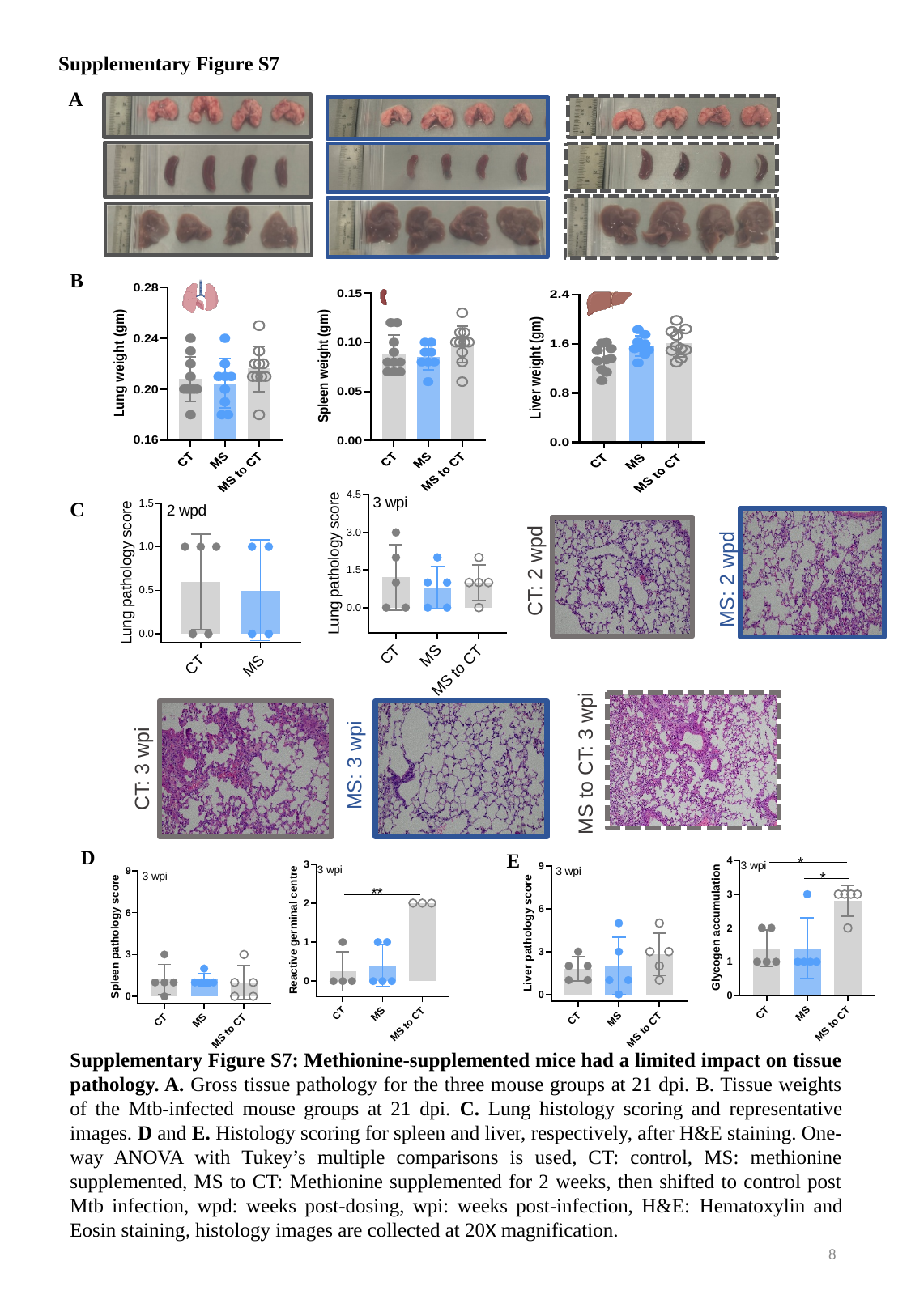

Supplementary Figure S7
A
B
MS: 2 wpd
CT: 2 wpd
C
MS to CT: 3 wpi
MS: 3 wpi
CT: 3 wpi
D
E
Supplementary Figure S7: Methionine-supplemented mice had a limited impact on tissue pathology. A. Gross tissue pathology for the three mouse groups at 21 dpi. B. Tissue weights of the Mtb-infected mouse groups at 21 dpi. C. Lung histology scoring and representative images. D and E. Histology scoring for spleen and liver, respectively, after H&E staining. One-way ANOVA with Tukey’s multiple comparisons is used, CT: control, MS: methionine supplemented, MS to CT: Methionine supplemented for 2 weeks, then shifted to control post Mtb infection, wpd: weeks post-dosing, wpi: weeks post-infection, H&E: Hematoxylin and Eosin staining, histology images are collected at 20X magnification.
8

### Slide 9
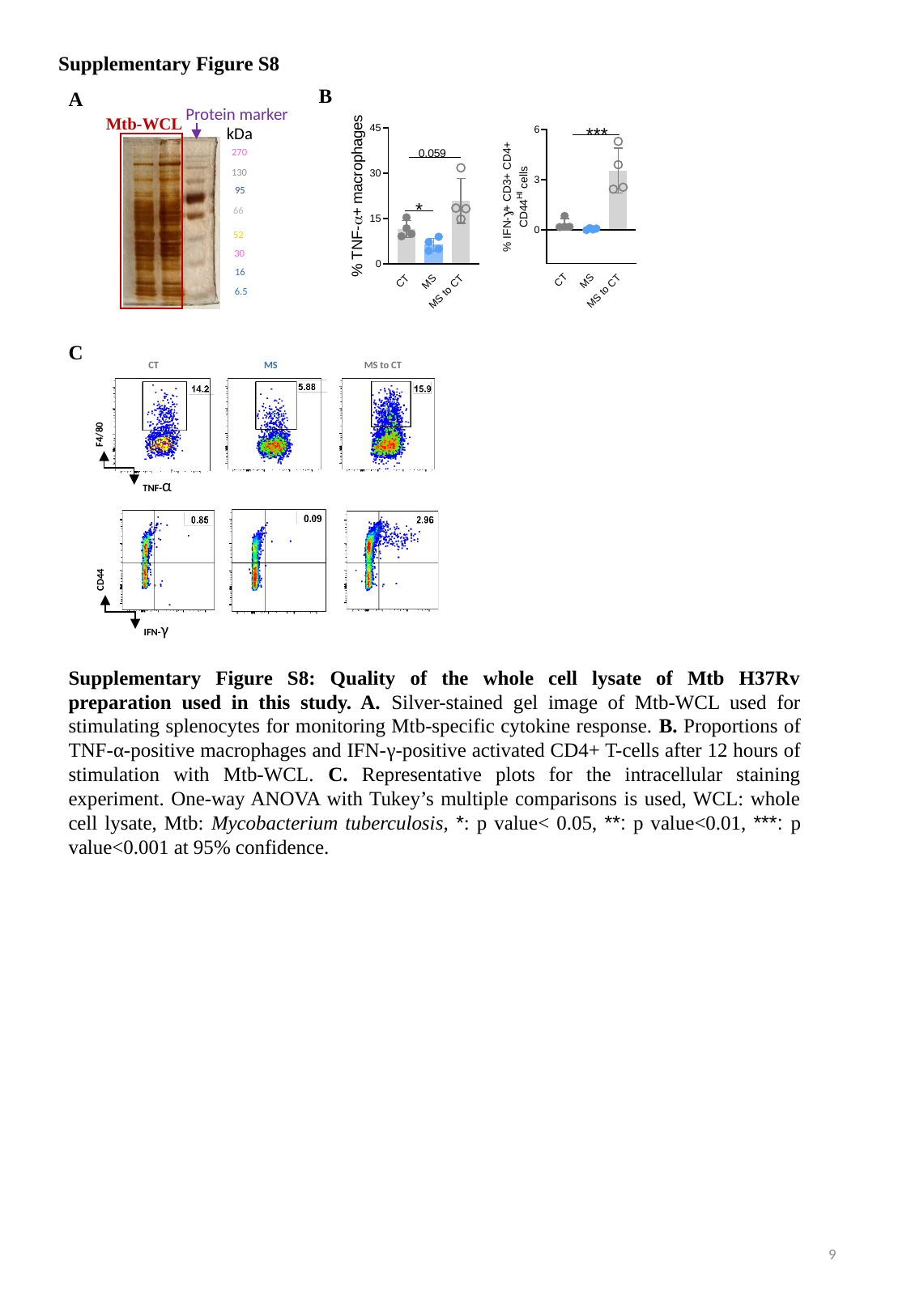

Supplementary Figure S8
B
A
Protein marker
Mtb-WCL
270
130
95
66
52
30
16
6.5
kDa
C
MS to CT
CT
MS
F4/80
TNF-α
CD44
IFN-γ
Supplementary Figure S8: Quality of the whole cell lysate of Mtb H37Rv preparation used in this study. A. Silver-stained gel image of Mtb-WCL used for stimulating splenocytes for monitoring Mtb-specific cytokine response. B. Proportions of TNF-α-positive macrophages and IFN-γ-positive activated CD4+ T-cells after 12 hours of stimulation with Mtb-WCL. C. Representative plots for the intracellular staining experiment. One-way ANOVA with Tukey’s multiple comparisons is used, WCL: whole cell lysate, Mtb: Mycobacterium tuberculosis, *: p value< 0.05, **: p value<0.01, ***: p value<0.001 at 95% confidence.
9

### Slide 10
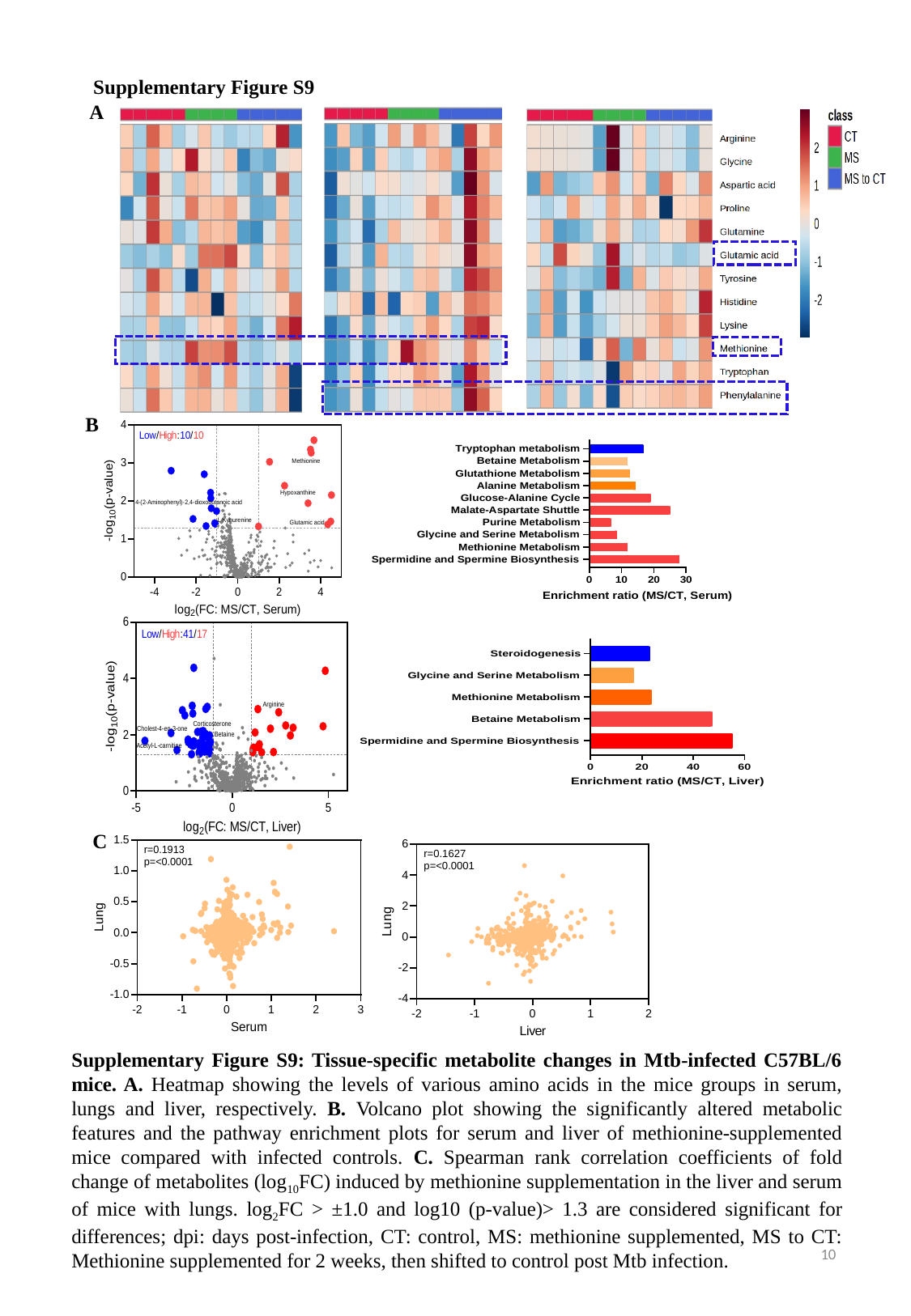

Supplementary Figure S9
A
B
C
Supplementary Figure S9: Tissue-specific metabolite changes in Mtb-infected C57BL/6 mice. A. Heatmap showing the levels of various amino acids in the mice groups in serum, lungs and liver, respectively. B. Volcano plot showing the significantly altered metabolic features and the pathway enrichment plots for serum and liver of methionine-supplemented mice compared with infected controls. C. Spearman rank correlation coefficients of fold change of metabolites (log_10_FC) induced by methionine supplementation in the liver and serum of mice with lungs. log_2_FC > ±1.0 and log10 (p-value)> 1.3 are considered significant for differences; dpi: days post-infection, CT: control, MS: methionine supplemented, MS to CT: Methionine supplemented for 2 weeks, then shifted to control post Mtb infection.
10

### Slide 11
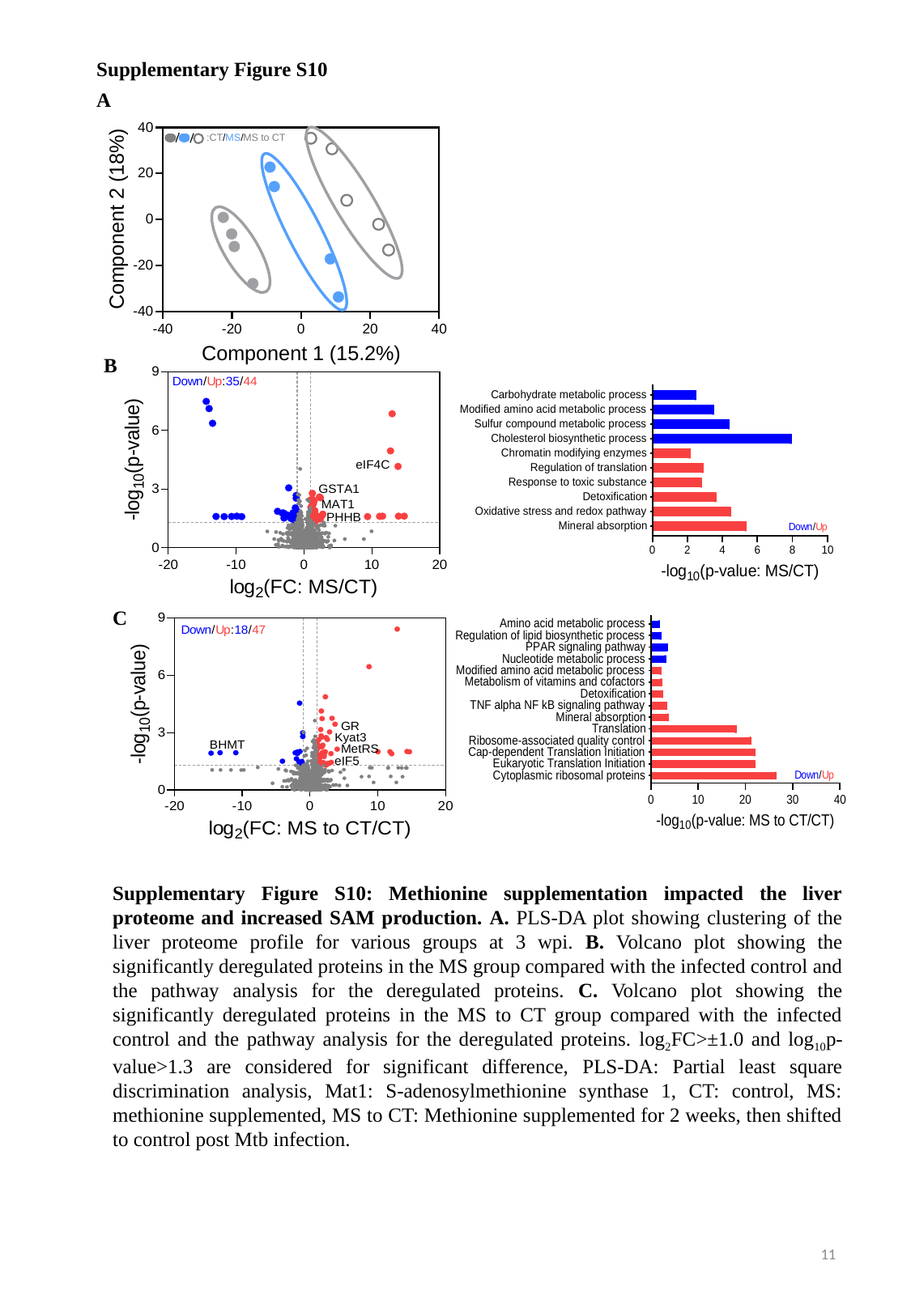

Supplementary Figure S10
A
B
C
Supplementary Figure S10: Methionine supplementation impacted the liver proteome and increased SAM production. A. PLS-DA plot showing clustering of the liver proteome profile for various groups at 3 wpi. B. Volcano plot showing the significantly deregulated proteins in the MS group compared with the infected control and the pathway analysis for the deregulated proteins. C. Volcano plot showing the significantly deregulated proteins in the MS to CT group compared with the infected control and the pathway analysis for the deregulated proteins. log_2_FC>±1.0 and log_10_p-value>1.3 are considered for significant difference, PLS-DA: Partial least square discrimination analysis, Mat1: S-adenosylmethionine synthase 1, CT: control, MS: methionine supplemented, MS to CT: Methionine supplemented for 2 weeks, then shifted to control post Mtb infection.
11

### Slide 12
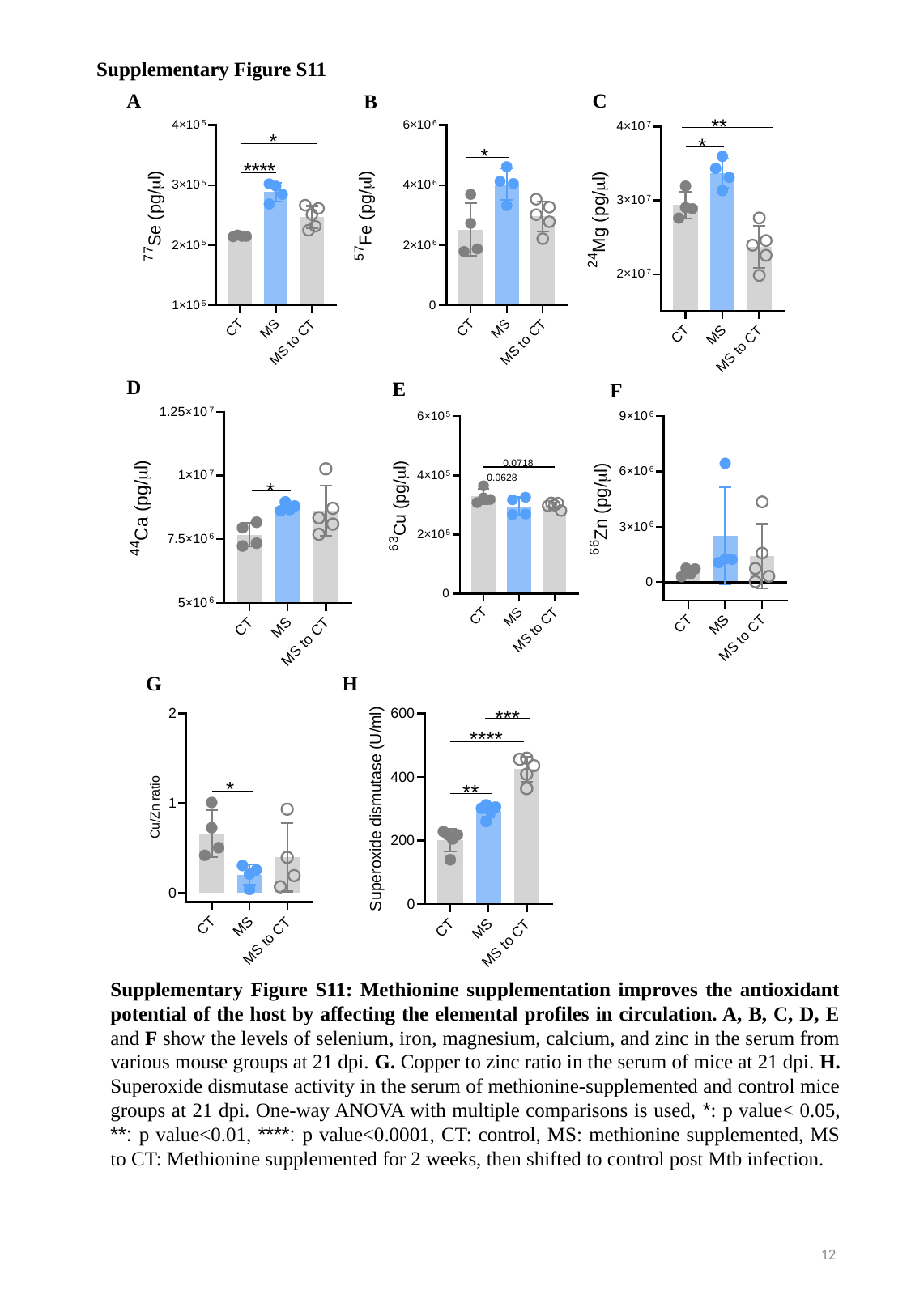

Supplementary Figure S11
A
C
B
D
E
F
G
H
Supplementary Figure S11: Methionine supplementation improves the antioxidant potential of the host by affecting the elemental profiles in circulation. A, B, C, D, E and F show the levels of selenium, iron, magnesium, calcium, and zinc in the serum from various mouse groups at 21 dpi. G. Copper to zinc ratio in the serum of mice at 21 dpi. H. Superoxide dismutase activity in the serum of methionine-supplemented and control mice groups at 21 dpi. One-way ANOVA with multiple comparisons is used, *: p value< 0.05, **: p value<0.01, ****: p value<0.0001, CT: control, MS: methionine supplemented, MS to CT: Methionine supplemented for 2 weeks, then shifted to control post Mtb infection.
12

### Slide 13
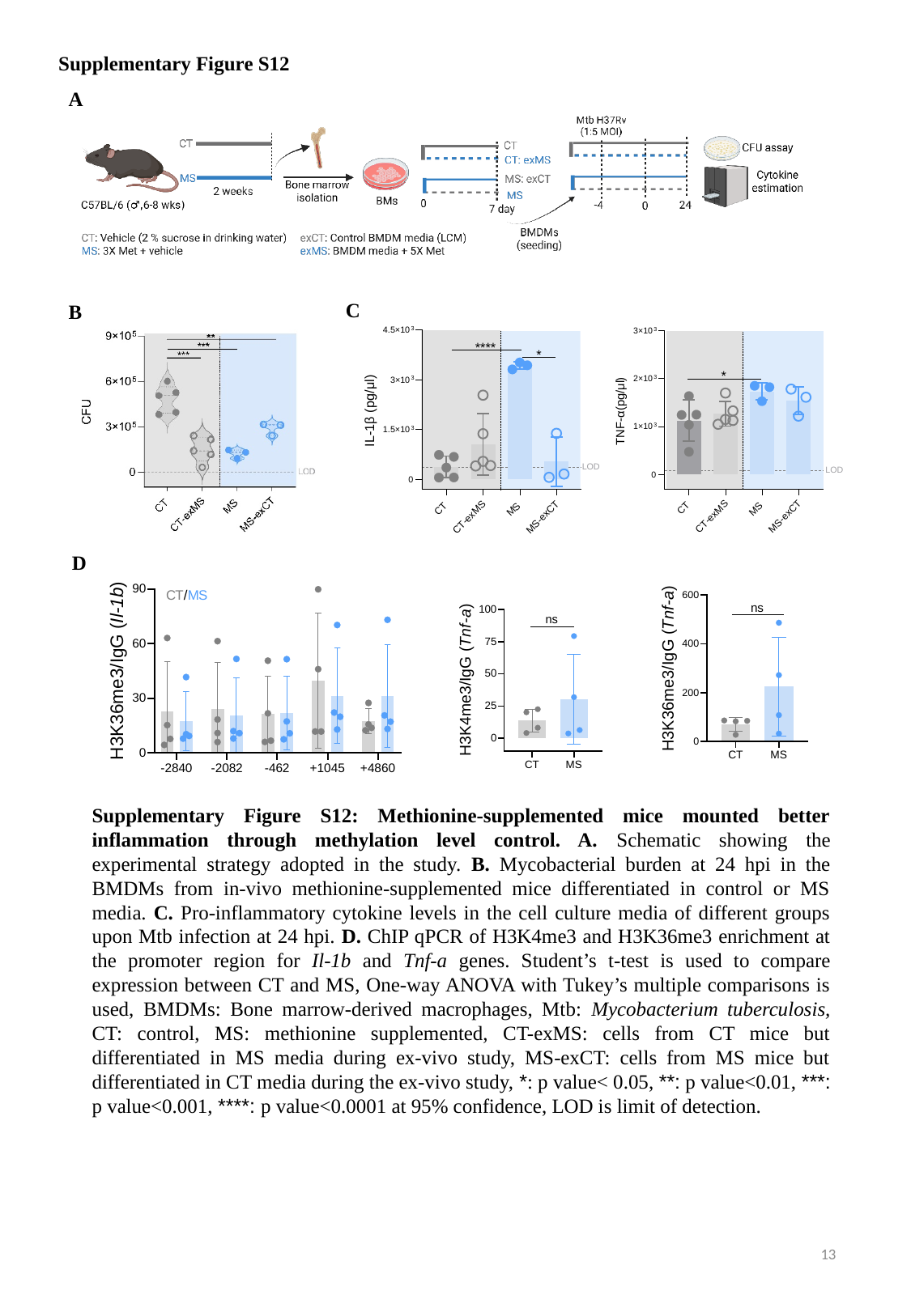

Supplementary Figure S12
A
C
B
D
Supplementary Figure S12: Methionine-supplemented mice mounted better inflammation through methylation level control. A. Schematic showing the experimental strategy adopted in the study. B. Mycobacterial burden at 24 hpi in the BMDMs from in-vivo methionine-supplemented mice differentiated in control or MS media. C. Pro-inflammatory cytokine levels in the cell culture media of different groups upon Mtb infection at 24 hpi. D. ChIP qPCR of H3K4me3 and H3K36me3 enrichment at the promoter region for Il-1b and Tnf-a genes. Student’s t-test is used to compare expression between CT and MS, One-way ANOVA with Tukey’s multiple comparisons is used, BMDMs: Bone marrow-derived macrophages, Mtb: Mycobacterium tuberculosis, CT: control, MS: methionine supplemented, CT-exMS: cells from CT mice but differentiated in MS media during ex-vivo study, MS-exCT: cells from MS mice but differentiated in CT media during the ex-vivo study, *: p value< 0.05, **: p value<0.01, ***: p value<0.001, ****: p value<0.0001 at 95% confidence, LOD is limit of detection.
13
